# Linking rhizosphere processes across scales: Opinion

**DOI:** 10.1101/2021.07.08.451655

**Authors:** A. Schnepf, A. Carminati, M. A. Ahmed, M. Ani, P. Benard, J. Bentz, M. Bonkowski, M. Brax, D. Diehl, P. Duddek, E. Kröner, M. Javaux, M. Landl, E. Lehndorff, E. Lippold, A. Lieu, C. W. Mueller, E. Oburger, W. Otten, X. Portell, M. Phalempin, A. Prechtel, R. Schulz, J. Vanderborght, D. Vetterlein

**Affiliations:** Forschungszentrum Jülich GmbH, IBG-3 (Agrosphäre), Wilhelm Johnen Str., D-52428 Jülich, Germany; ETH Zürich, Department of Environmental Systems Science, Universitätstr. 16, 8092 Zürich, Switzerland; Chair of Soil Physics, Bayreuth Center of Ecology and Environmental Research (BayCEER), University of Bayreuth, Universitätsstr 30, 95447, Bayreuth, Germany; University of Koblenz-Landau, Institute for Environmental Sciences, Fortstr. 7, 76829 Landau, Germany; University of Cologne, Institute of Zoology, Zülpicher Str. 47b, 50674 Cologne, Germany; University Bonn, Bonn, Institute of Crop Science and Resource Conservation (INRES), Karlrobert-Kreiten-Straße 13, 53115 Bonn, Germany; Université catholique de Louvain, Earth and Life Institute, Croix du Sud L7.05.02, B-1348 Louvain-la-Neuve, Belgium; Department of Soil System Science, Helmholtz Centre for Environmental Research – UFZ, Halle, Germany; Friedrich-Alexander University of Erlangen-Nürnberg, Department of Mathematics, Cauerstr. 11, 91058 Erlangen, Germany; University of Copenhagen, Department of Geosciences and Natural Resource Management, Øster Voldgade 10, 1350 Copenhagen K, Denmark; University of Natural Resources and Life Sciences, Institute of Soil Research, Konrad Lorenz-Str. 24, 3430 Tulln an der Donau, Austria; School of Water, Energy and Environment, Cranfield University, Cranfield, Bedfordshire MK43 0AL, UK, Cranfield University, UK

**Keywords:** rhizosphere, modelling, up- and downscaling, emergent behaviour

## Abstract

**Purpose:** Simultaneously interacting small-scale rhizosphere processes determine emergent plant-scale behaviour, including growth, transpiration, nutrient uptake, soil carbon storage and transformation by microorganisms. Current advances in modelling and experimental methods open the path to unravel and link those processes.

**Methods:** We present a series of examples of state-of-the art simulations addressing this multi-scale, multi-process problem from a modelling point of view, as well as from the point of view of integrating newly available rhizosphere data and images.

**Results:** Each example includes a model that links scales and experimental data to set-up simulations that explain and predict spatial and temporal distribution of rhizodeposition as driven by root architecture development, soil structure, presence of root hairs, soil water content and distribution of soil water. Furthermore, two models explicitly simulate the impact of the rhizodeposits on plant nutrient uptake and soil microbial activity, respectively. This exemplifies the currently available state of the art modelling tools in this field: image-based modelling, pore-scale modelling, continuum scale modelling and functional-structural plant modelling. We further show how to link the pore scale to the continuum scale by homogenisation or by deriving effective physical parameters like viscosity from nano-scale chemical properties.

**Conclusion:** Modelling allows to integrate and make use of new experimental data across different rhizosphere processes (and thus across different disciplines) and scales. Described models are tools to test hypotheses and consequently improve our mechanistic understanding of how rhizosphere processes impact plant-scale behaviour. Linking multiple scales and processes is the logical next step for future research.

## Introduction

The rhizosphere is one of the most complex and vital interfaces on earth. It hosts myriads of microorganisms, and its properties affect terrestrial fluxes of water and various elements including carbon and nitrogen. Rhizosphere properties are the result of manifold biological, physical, and chemical processes that ultimately impact plant growth and soil properties. These processes include water and nutrient uptake, rhizodeposition and microbial activity, and rearrangement of soil particles by the root as it grows. These processes *interactively* affect each other and dynamically determine the rhizosphere properties.

The objective of this opinion paper is to propose a road map for integrating detailed rhizosphere processes into a plant-scale and soil profile-scale modelling concept. The working hypothesis of the proposed approach is that the emergent behaviour at the plant scale is determined by the combined **interaction** of small-scale rhizosphere processes (Vetterlein et al., 2020). The emergent behaviour includes plant growth (e.g. biomass), bulk soil properties (e.g. infiltrability), transpiration, carbon fluxes, nutrient uptake, plant health, soil aggregation, and carbon storage and transformation. Small-scale rhizosphere processes include root exudation, microbial transformation and biodiversity, water and nutrient uptake.

We further distinguish between *effective properties*, which result from upscaling small-scale properties to larger scales, and *emergent behaviour*, which develops from the interactions between multiple small-scale processes.

On the one hand, our approach is to combine time-series images down to the pore scale with detailed models to define effective properties to be included in plant-scale architecture models. On the other hand, our approach proposes to use large-scale models and information to feed into the small-scale models addressing longer temporal scales. Doing so, we propose a cross-talk between detailed and emerging processes, with both up- and down-scaling. A sketch of the scales and related processes we aim to address is shown in Fig. 1. The objective of *upscaling* multiple small-scale processes is to *determine effective properties and emergent behaviours* and to predict qualitative behaviours and trends. The objective of *downscaling* is to *set boundary conditions* for small-scale models which are used to resolve smaller scale structures and evaluate their effect on emergent behaviours at larger scales.

**Fig. 1.**
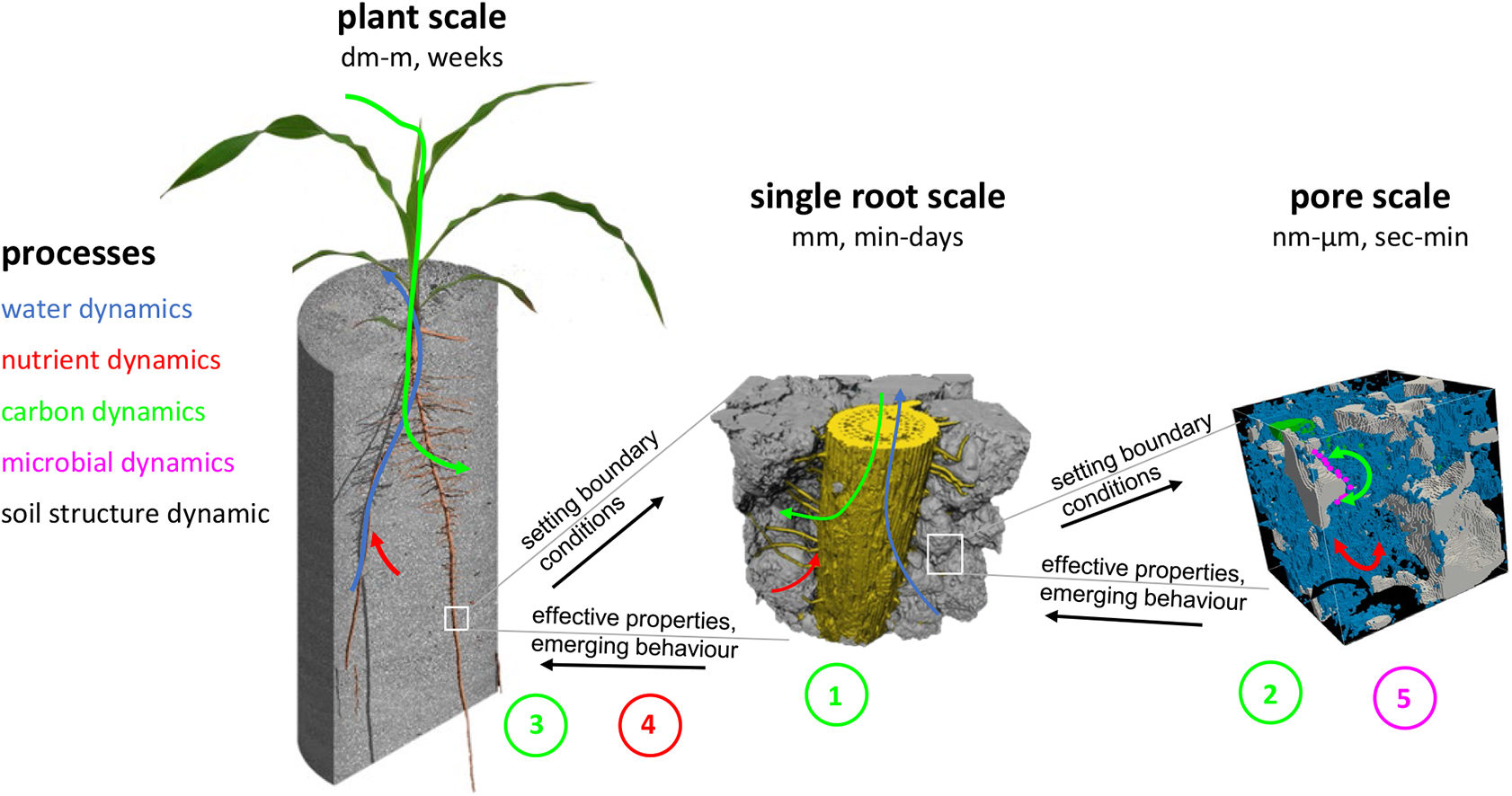
Linking rhizosphere processes across scales as illustrated by the examples 1-5 presented in this opinion paper. Colours of arrows and examples correspond to the different processes mentioned on the left.

### Examples of linking mechanistic models and data across scales

In the following paragraphs, we present a series of examples of state-of-the art modelling approaches that involve upscaling from single root to plant scale or from pore scale to single root scale, and we discuss current challenges involved in considering multiple scales and processes in one simulation. Some of the examples are based on published data sets while others use yet unpublished works that are intended to serve as showcases. The examples address how existing data could be integrated into the respective models.

Even if in this contribution we will not explicitly combine all possible processes (just some as examples), we would like to highlight that all these processes occur on multiple scales and are indeed linked.

All examples use the same basic structure: question, scales, approach, results, challenges, and open questions. *Example 1* is an example of image-based modelling of the diffusion of root exudates within a soil domain that was captured and meshed from a micro computed tomography (µCT) image at the pore scale. *Example 2* also used µCT images of soil at the pore scale to derive the effective diffusion coefficient at the continuum scale (here used for both the single root and plant scales) for scenarios with different mucilage and water contents and distributions. This example also incorporates data about physico-chemical properties of mucilage. *Example 3* couples a 3D dynamic root architecture simulation with root exudation of individual roots to compute the rhizodeposition patterns at the plant scale as driven by multiple single root growth and rhizodeposition properties. *Example 4* quantifies the impact of root exudation by multiple roots of a plant to root phosphorus uptake. The overall plant phosphorus uptake and carbon investment emerge from these single root interactions. *Example 5* uses an explicit description of the 3D soil structure, water, and carbon distributions at the pore scale along with an individual-based approach describing microbial dynamics. In this approach, microbial driven processes emerge from interactions among the biotic and abiotic components described.

### Example 1: Root exudation and the role of root hairs on the single root scale

**Question** How do root hairs and the contact between the root surface and the soil matrix affect rhizodeposition and the spatial distribution of root exudates from the root surface?

**Scales** Spatial scale: <1 mm, temporal scale: < 1 day

### Approach

#### Data Acquisition and Processing

Maize plants (*Zea mays L*.) were grown in seedling holder microcosms (Keyes et al., 2013; Koebernick et al., 2017) which were filled with a sieved and fertilized loamy substrate (Vetterlein et al., 2021). For image acquisition, roots of the 14 days old plants were scanned non-destructively at an actual pixel size of 0.65µm using a synchrotron radiation X-ray CT (TOMCAT, Paul Scherrer Institute Villigen, Switzerland). In order to segment the scanned root compartments as well as soil aggregates, image processing was performed in Avizo. 3D finite element meshes consisting of the soil matrix and the root-soil contact were generated in Gmsh.

#### Modelling

To assess the effect of root hairs on the spatial distribution of root exudates, image-based modelling on the pre-processed CT data was performed. Diffusion simulations of carbon released by a root compartment of approx. 1.4 mm length into the rhizosphere were carried out for two cases on one exemplary sample – a root compartment with and without hairs. We considered the root as well as its’ hairs to release carbon (Holz et al., 2017) treating their contact surface to soil aggregates as inlet at a constant concentration. Carbon diffusion was calculated in the partially saturated soil micropore region with a no-flow boundary condition at the surface of the soil aggregates. There was no carbon diffusion in the air-filled macropores and, furthermore, we neglected both root and root hair growth.

Mathematically, the underlying problem is described by the diffusion equation - a second order parabolic partial differential equation:

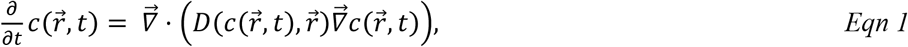

where 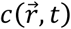 denotes the concentration of the diffusing material at location 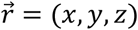 and time *t*. 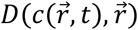 represents the diffusion coefficient for a concentration *c* at a location 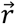 and 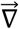 the nabla operator.

Assuming a constant diffusion coefficient, our model is formulated as follows:

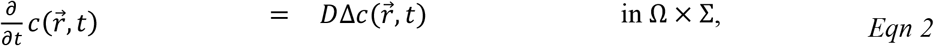

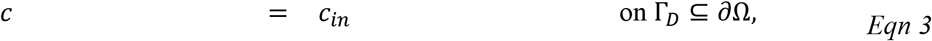

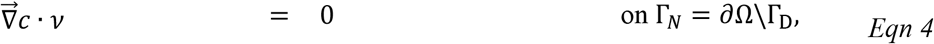

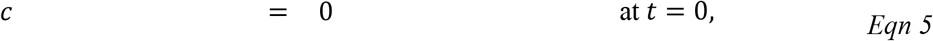

With

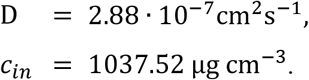

The domain obtained from CT images is denoted by Ω, Σ = [0, 3600*s*] is the simulated time interval. *v* represents the unit outer normal and Δ the Laplacian operator. The inlet carbon concentration *c*_*in*_ was taken to be a fixed value and taken from Holz et al. (2018) and the diffusion coefficient *D* is 3.3 times lower but still comparable to the one obtained for glucose in soil using the equation proposed by Millington and Quirk (1961).

The diffusion equation was discretized and solved by the “scalarTransportFoam” solver of OpenFoam – an open source CFD software package (Weller et al., 1998). Further details are provided in the supplementary information.

Fig. 2 outlines the workflow of example 1. Data acquisition includes conducting an experiment of plant growth in a soil column and the imaging of this soil column in a synchrotron facility. The raw images are processes in a way that results in a 3D finite element mesh of the soil domain on which diffusion of root-derived exudates was performed using image-based modelling techniques. The outcome is the spatio-temporal pattern of exudate concentration on the pore-scale.

**Fig. 2.**
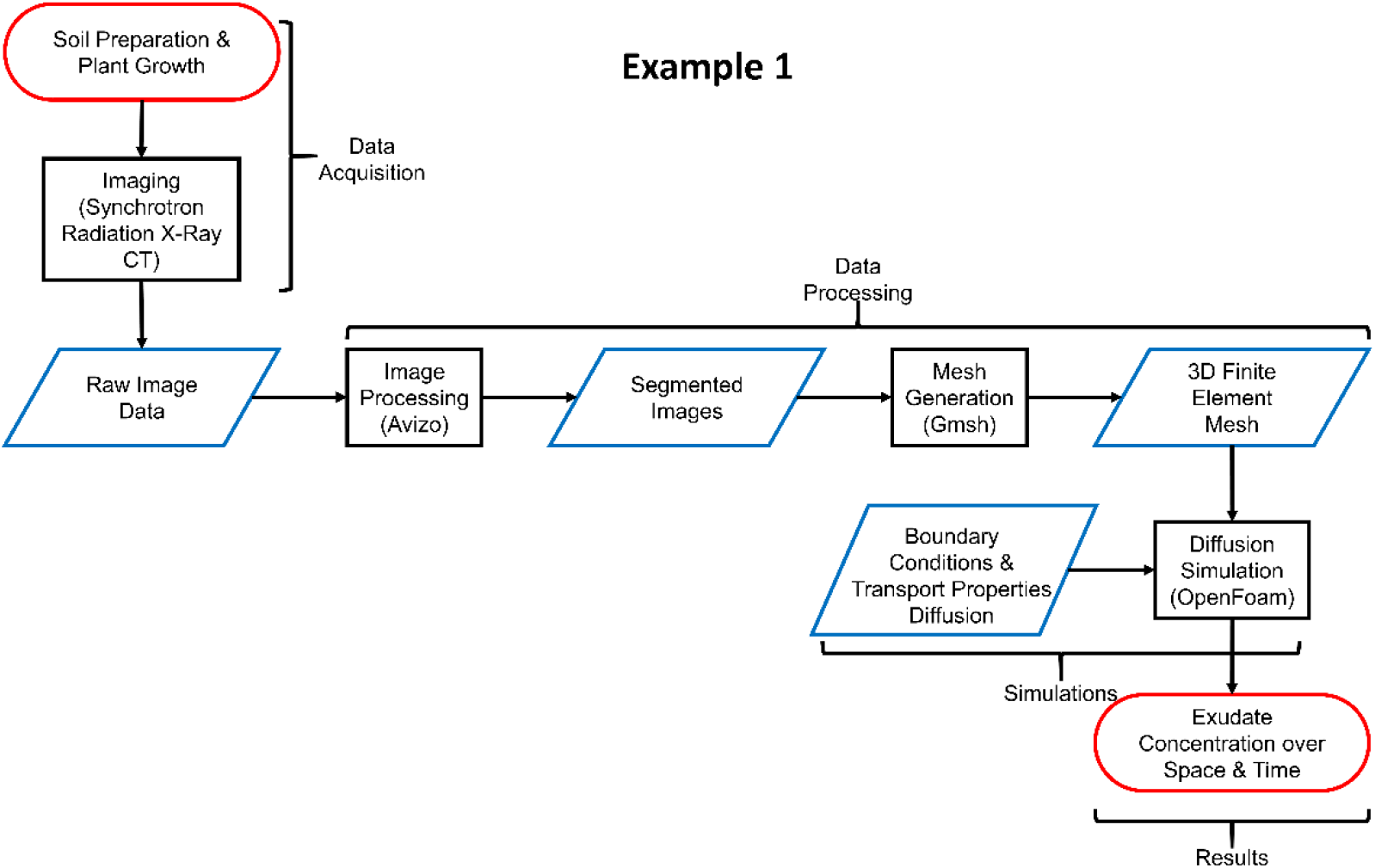
Flow chart outlining the approach of example 1

## Results

We simulated the diffusion of root exudates, in particular carbon, into soil for one hour and one illustrative sample. The selected region of interest of approx. 1.8 mm x 1.3 mm x 1.4 mm contained a soil aggregate volume of 1.3 mm^3^, an air-filled macropore volume of 1.1 mm^3^, a root volume of 0.4 mm^3^ and an air volume of 0.4 mm^3^that surrounded the sample tube. The epidermis surface area was 3.9 mm^2^ and a fraction of 6.1% was in contact to soil. The soil contact fraction of the root including hairs was approximately three times bigger (17.7%).

Regarding the hairless root, a total carbon mass of 0.067 µg diffused into the soil after 1 hour, whereas this value was three times higher for the root with hair (0.204 µg). The comparison of carbon distributions within the rhizosphere revealed a lower radial concentration gradient for the hair case resulting in a right shift of the concentration data (Fig. 3c). In the hairless case, carbon concentration dropped below 1% at a distance of 0.29 mm from the root surface whereas the same value was reached in the hair-case at 0.74 mm.

**Fig. 3.**
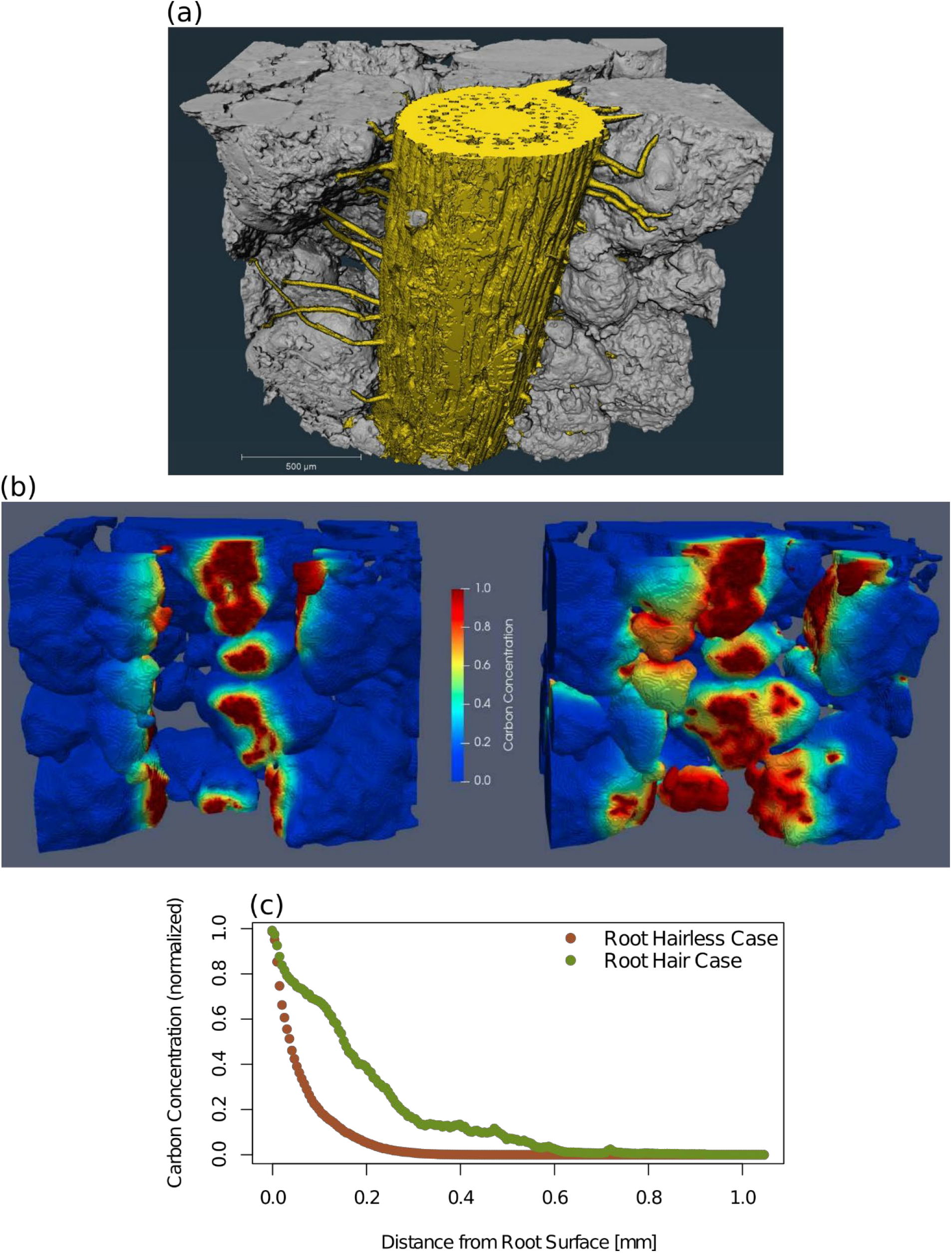
Results of image analysis and simulations. (a) 3D rendering of the segmented synchrotron X-ray CT sample consisting of the root with its elongated hairs in yellow and soil particles in grey; scale bar = 500 µm. (b) Illustrative comparison of the simulated carbon diffusion within the soil domain for the hairless root (left) and the root with hair (right) after a simulation time of 1 hour. (c) Spatial carbon distribution within the rhizosphere represented by the carbon concentration regarding the radial distance from the root-surface.

### Challenges and open questions

This illustrative example shows how root hairs can increase the total amount of root exudates and their diffusional distance into the soil if the exudation rate is constant. Explicit image-based simulations of root exudation including information of root-soil contact and the spatial distribution of root hairs allows estimating the importance of root hairs for rhizodeposition. This example shows good agreement with experimental measurements by Holz et al. (2017).

Note that we did not compare a wildtype with a root hair defective mutant and we assumed the same input concentration for both genotypes. Our simulations are based on the grid of a single wildtype sample before and after removing its’ root hair domain. Furthermore, the focus of this example lies on exudate diffusion from a static (non-growing) root.

One open question is how to implicitly account for hairs when their spatial distribution as well as the contacts with the soil are not known and how to include such implementations in root system models. The outcome of our simulation may be used to define the boundary conditions of the simulations performed in example 5. An additional open question is the representativity of the selected volume. Indeed, soil porosity and root hair density might be highly variable in space (at the scale of 1 mm^3^) and time. Temporal information is needed to cover the lifetime of root hairs. Additionally, rhizosphere bacteria were shown to contribute to mucilage production, forming jointly with root mucilage a rhizosheath in maize whose extend is determined by soil moisture (Watt et al., 1994; Watt et al., 1993). Therefore different soil water contents and soil textures should be simulated in order to have a representative picture of the role of root hairs on rhizodeposition for variable conditions. Finally, we did not consider microbial degradation of exudates, which is discussed in example 5.

In summary, the biggest challenges are related to the representativity of the imaged soil volume and how to integrate these simulations in models that consider root growth and other interacting processes, such as root water uptake (affecting soil water and the contacts) and microbial degradation of the rhizodeposits.

### Example 2: Mucilage and hydraulic properties

**Question** How do the intrinsic chemical properties of mucilage affect pore scale distribution and dynamics of water and mucilage during drying? How does the altered liquid configuration affect the rhizosphere effective hydraulic properties and solute diffusion?

**Scales** Spatial scale: from molecular to nm, µm and to mm, temporal scale: < 1 day

### Approach

#### Background

Root mucilage is primarily released at root tips and mainly composed of polysaccharides, proteins and some lipids. In contact with water, these polymers swell and form a 3D gel network. Gels possess specific properties, such as water holding capacity, the ability of swelling and shrinking, and viscoelasticity, which impact soil functions, such as water retention (Ahmed et al., 2014; Kroener et al., 2018) and nutrient supply (Brax et al., 2017).

In contrast to pure water whose distribution in soil is primarily controlled by its surface tension and capillary forces, the liquid configuration of mucilage is additionally determined by its chemical hydrogel properties and their physical stability which rely on structure and arrangement of the contained polysaccharide units.

Gel properties vary for mucilage located in soil pores in contrast to “free” mucilage (Brax et al., 2020; Brax et al., 2019). The polymers grip to the soil particle surface and the network has thus an increased strength compared to the “free” gel. The confinement by the pore walls leads to an inhomogeneous distribution of the polymer network in the pore during swelling and shrinking (Marcombe et al., 2010) and to pore-size specific organization of the polymeric network.

Upon drying, the concentration of polysaccharides within the liquid phase increases and the internal structural units of mucilage become more and more important in controlling the physical properties of the liquid solution and pore scale distribution of the liquid or hydrogel phase. Thereby, mucilage goes through a glassy transition, passing from a liquid to a hydrogel and finally to a rather solid structure.

This leads to characteristic mucilage drying patterns that depend on intrinsic physical mucilage properties like viscosity and surface tension. In turn, the physical properties of mucilage depend on its chemical composition, structural stabilization and, thus, the supramolecular arrangement of the polymers. Note that chemical properties of mucilage vary with plant species (Brax et al., 2020) and environmental conditions (pH, ionic strength, cations, surfactants). For instance, viscosity of maize root mucilage is higher than that of wheat.

The different viscosities of wheat and maize mucilage explain different patterns formed at the nano-scale by the respective mucilage upon drying on a flat surface (Fig. 4a,b). For wheat, the nanostructure appears as a network characterized by thin branches smaller than 30 nm in width. Maize mucilage builds a more connected coating with larger hole-structures. The thin threads of wheat mucilage suggest less and weaker interactions between polymers compared to maize mucilage, which is consistent with the differences in viscosity (Fig. 4c).

**Fig. 4.**
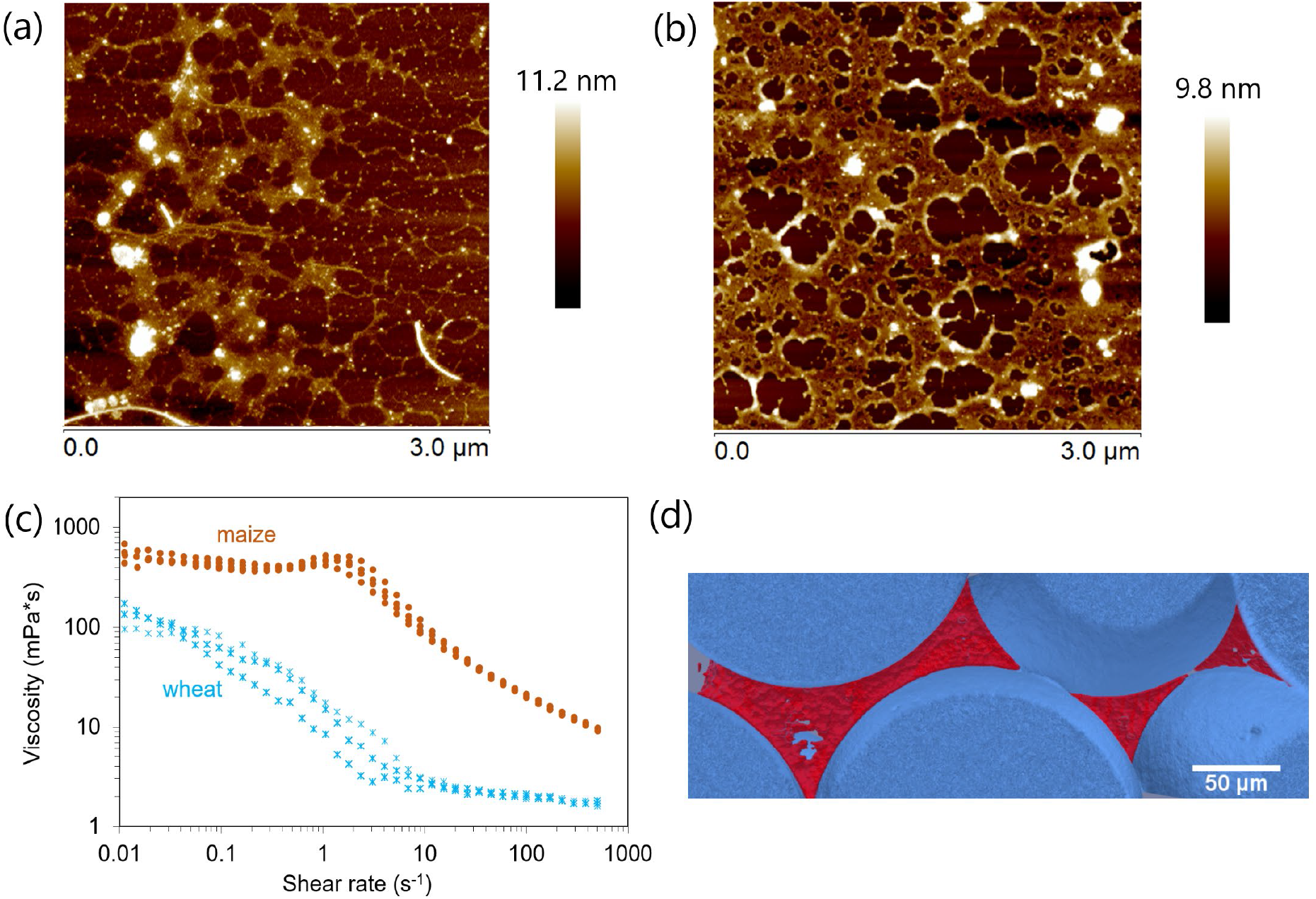
(a) Atomic force microscopy height image of (a) wheat and (b) maize root mucilage dried on flat mineral surface taken in Peak-Force Quantitative Nanoscale Mechanical (PFQNM) mode; (c) flow curves of viscosity at various shear rates measured for maize and wheat root mucilage at 7 mg/mL with three replicates for each species; (d) mucilage from maize nodal roots let dry in glass beads (100-200 µm in diameter) imaged via X-ray CT. The content of mucilage was 8 mg g^-1^ (weight of dry mucilage per weight of dry particles). An interconnected 2D surface of dry mucilage was deposited through multiple pores upon drying (Benard et al., 2019).

Analogue pictures are visible in 3D porous media. Figure 4d shows that upon drying in a porous medium mucilage forms an interconnected surface that spans through multiple pores, maintaining the liquid phase connected during the drying period. Such a configuration of the liquid phase upon severe drying is very distinct from that of water, whose high surface tension causes the breakup of liquid bridges. For mucilage, other forces come into play, which include the resistance of water to flow through the polymer network and the forces needed to deform and stretch the polymer network.

#### Spatial configuration

To simulate the interplay between the molecules of the polymer network and water, modelling tools are needed that can describe both, liquid dynamics and dynamics of the polymeric network. Lattice-Boltzmann methods are common tools to simulate pore scale dynamics of liquids (Pot et al., 2015; Richefeu et al., 2016; Sukop and Or, 2004; Tuller and Or, 2005). Discrete element methods are tools to describe deformation and rupture processes of solids (Bobet et al., 2009) and have been used to simulate fracture of hydrogels (Kimber et al., 2012; Yang et al., 2018). While most previous hydrogel simulations consider free hydrogels, here, we study hydrogel deformation when it is confined within pore spac **Challenges and open questions** e and attached to soil particle surfaces.

#### Transport properties

Both, phase distributions and connectivities as well as intrinsic mucilage properties will finally affect hydraulic properties, gas diffusion and nutrient transport. Macroscale models cannot take into account the explicit geometries and properties of phase distributions at the pore scale. The pore scale model however is not amenable to large-scale computations because of its high complexity. Information from the microscale can be incorporated to the single root scale using mathematical upscaling or homogenisation techniques (Ray et al., 2018) These methods allow, e.g., for the computation of, potentially anisotropic, effective diffusion coefficient tensors requiring only the geometric information within a representative elementary volume. These methods have been used, e.g., for nutrient diffusion with regular geometries (Leitner et al., 2010b), but have been extended recently also to irregular, evolving structures (Ray et al., 2018). Although the setting is periodic at the boundaries of the investigated domain, the underlying geometries can be arbitrarily complex. Using CT-scans of maize root mucilage distributions and realistic liquid configurations, we can study the impact of the presence of mucilage at the microscale on the effective diffusion coefficient – which can reduce or enhance the effective diffusion coefficient depending on the pore scale configuration and water (and mucilage) content.

Fig. 5 outlines the workflow of example 2. It is based on the same experimental setup as example 1, but uses additional modelling approaches to (a) simulate the water distribution at the pore scale for different water contents and mucilage concentrations using Lattice-Boltzmann and Discrete Element methods. Those amended images are then subject to mathematical homogenisation to compute effective diffusion tensors that may be used in continuum-scale simulations such as examples 3 and 4.

**Fig. 5.**
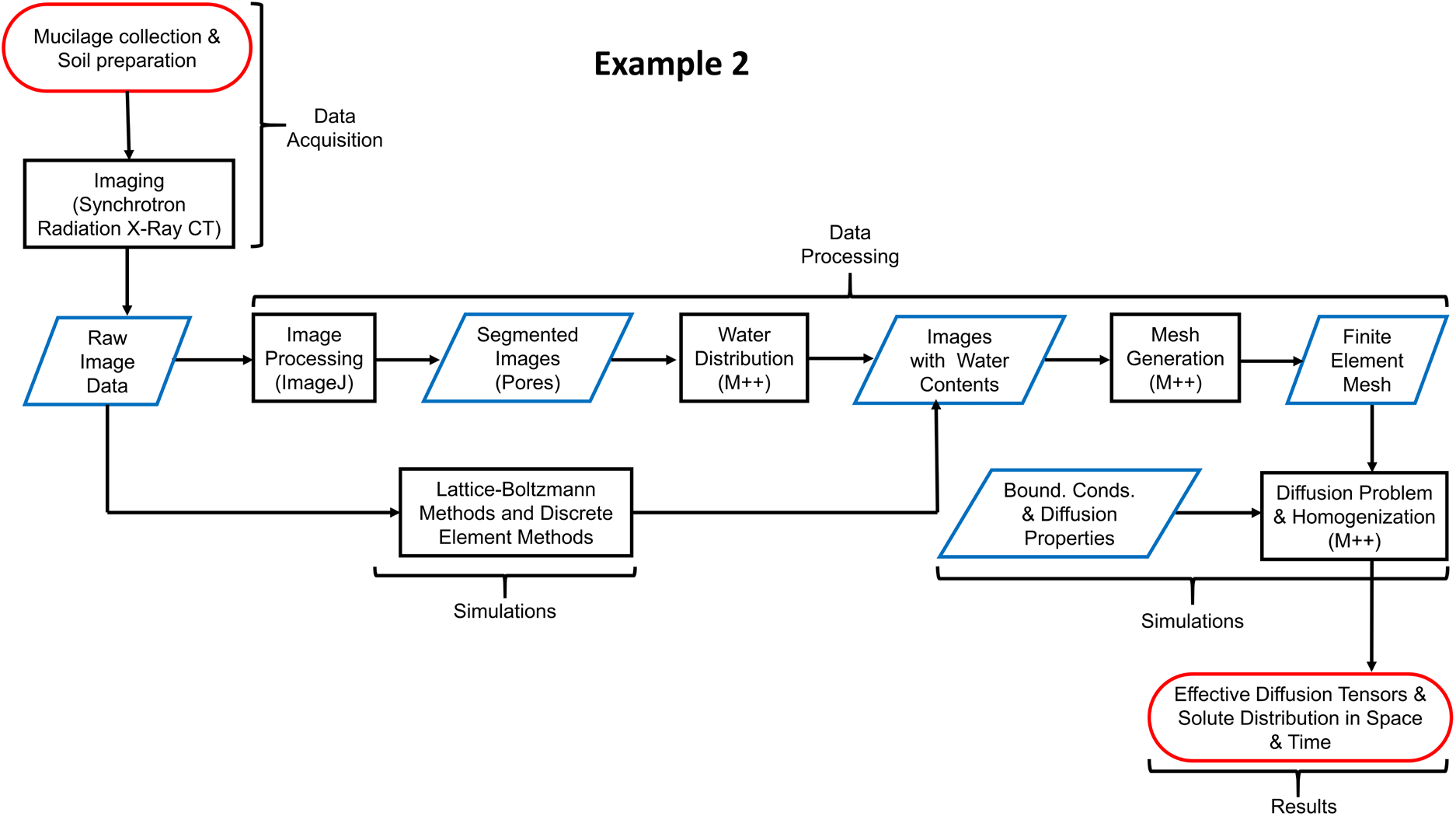
Flow chart outlining the approach of example 2.

## Results

### Spatial configuration

First simulations based on Lattice Boltzmann methods and discrete element methods, respectively, show the deformation of a liquid bridge between two soil particles upon drying (Fig. 6a). In the case of highly concentrated mucilage, some hollow structures can form that have also been observed in experiments (Benard et al., 2018). Due to these internal polymeric structural units mucilage bridges can still connect two particles under certain drying conditions at which water bridges would break. Similar results were obtained in a theoretical study (Carminati et al., 2017) where the increased viscosity of mucilage was held responsible for the damping effect on motion.

**Fig. 6.**
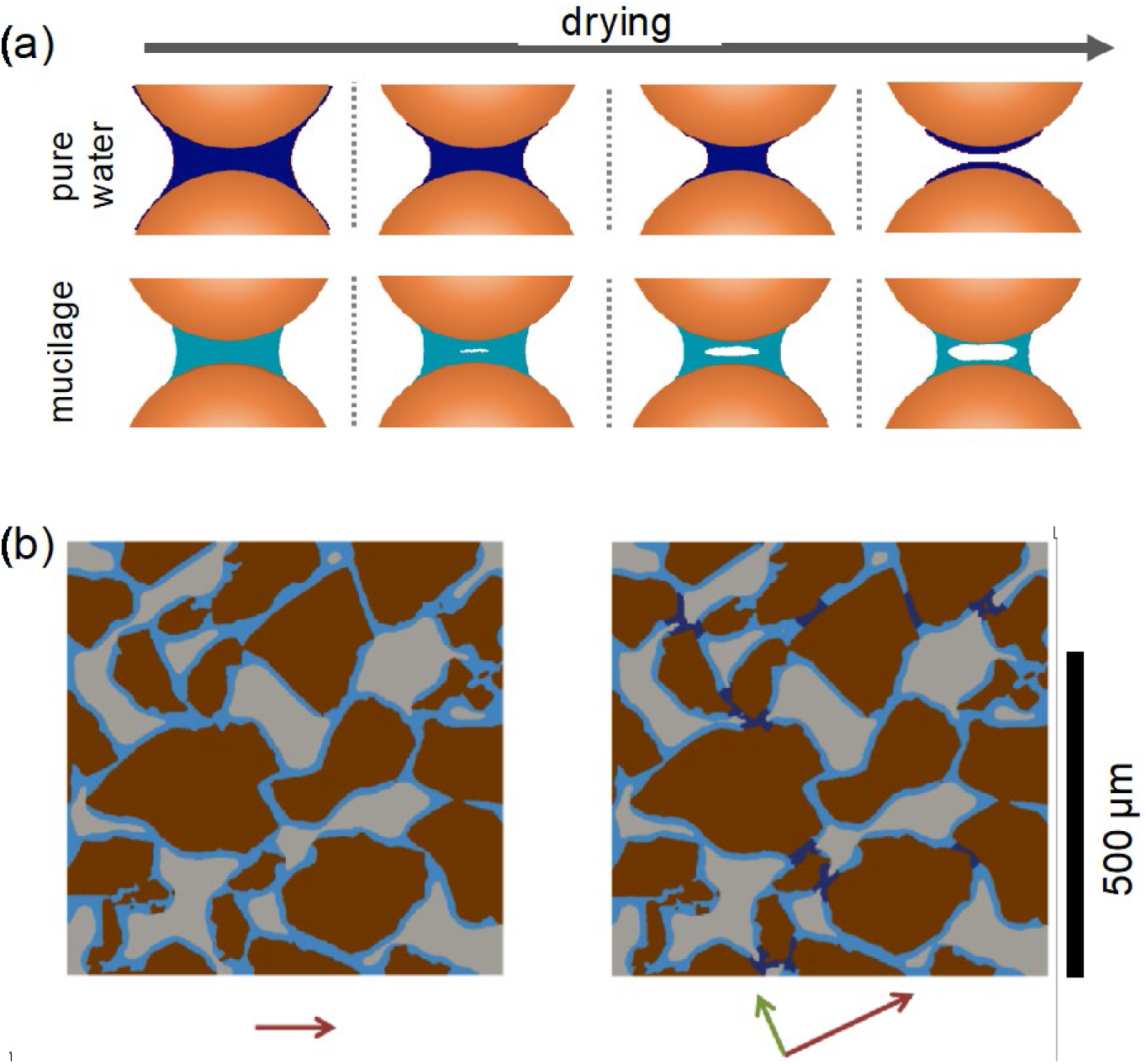
(a) Simulation of drying dynamics of a liquid bridge between two soil particles (modified from Haupenthal et al.). Top: pure water simulated using Lattice Boltzmann methods. Bottom: mucilage at high concentration simulated using the discrete element method; (b) 2D geometry derived from CT-scan [courtesy of P. Benard, M. Zarebanadkouki, University of Bayreuth, A. Carminati, ETH Zurich] of a sandy soil with porosity 41%. Artificial water distribution with water content of 19 % without (left) and with (right) mucilage. Sand particles (brown); air (grey); water (light blue); water and mucilage mixture (dark blue). The arrows represent the eigenvectors and eigenvalues (length), showing, in each case, the main directions of diffusion.

### Transport properties

Here we show how small changes in the spatial configuration of mucilage can affect the effective diffusion of solutes (e.g. nutrients) across the rhizosphere. We deal with the effect of mucilage on the effective diffusion coefficient, assuming that the presence of mucilage locally decreases the diffusion coefficient of a given chemical species. The species does not diffuse in the gas nor in the solid phases. The diffusion in water is set to a reference value of 1. In the mucilage, the diffusion is assumed to be Dmuc/Dref = 0.5, exemplarily.

We simulated a scenario in a sandy soil with a volumetric water content of 0.19 cm^3^cm^-3^. Figure 8b shows the geometries derived from a CT-scan used for two different scenarios. In the first scenario, the liquid phase consists of pure water only whereas the second includes pure water and liquid bridges induced by drying mucilage. In this particular case, the mucilage makes up only 10% of the liquid phase. The computed effective diffusion coefficients matrices now read:

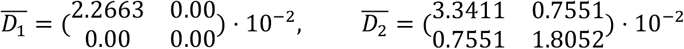

where 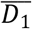 and 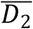 correspond to the matrices in the scenarios without and with mucilage respectively.

For both scenarios, the first diagonal entry is non-zero implying that the diffusion is achievable in the domain horizontally. However, in the other vertical direction, we notice that the diffusion is impeded though not impossible in the first scenario (≈ 10^−11^). Recall that the domain is periodic, i.e. the top boundary is continued with the lower boundary. So when a liquid cell on top meets a solid cell at the bottom, the pore is blocked there. In the second scenario the mucilage bridges keep the liquid phase more connected, creating new paths for the species to diffuse in the vertical direction. Note in particular the bridges (dark blue) at the top left and bottom left part of the right image in Figure 6b that connect previously disconnected areas. For each case, the main directions of diffusion are plotted in Figure 6. In this numerical experiment, the presence of mucilage affects the diffusion paths and hence increases the intensity of the diffusion. Although this is a 2D numerical study, it demonstrates that small differences in the connectivity of the phases between particles can imply large changes in effective diffusivity, and thus 3D pore scale information needs to be evaluated systematically in many samples to quantify the effect of mucilage on solute diffusion.

#### Challenges and open questions

We have demonstrated that the interaction between the polymer networks, water and soil particles increases the connectivity of the liquid phase across the rhizosphere. Upon drying, mucilage may be deposited as two-dimensional surfaces that may form hollow cylinders or interconnected surfaces across the pore space. Although the solute diffusion coefficient in mucilage is smaller than in pure water, its enhanced connectivity upon drying can result in an enhanced effective diffusion coefficient.

These physical mechanisms, which have been addressed in the models and the experimental observations, qualitatively match. However, several parameters and processes are still not yet considered in these models. To determine intrinsic physical mucilage properties, for example, it is necessary to consider nano-scale chemical properties such as content of high molecular weight material, the length and ramification of the polymers, the type of monomers and the resulting strength of interactions between the polymer chains. Also variations in the chemical environment like pH, absence or presence of mono- and multivalent cations or of organic surfactants may control the strength of the polymer network. These chemical properties need to be further investigated to predict how mucilage of various plants will affect biological and physical processes at the pore scale. Furthermore, it is not clear how to predict the typical mucilage content (gram of dry mucilage per soil dry weight) in the rhizosphere and to estimate at what concentration (gram of dry mucilage per liquid volume) the glassy transition takes place. For pore scale models, parameters of the Lattice-Boltzmann and the discrete element methods need to be related to measured physical properties of mucilage, such as viscosity, water adsorption, surface tension and elasticity of the polymeric structure. As soil texture, structure, and drying rate are likely to affect the spatial distribution of mucilage in soils, measurements of mucilage deposition for varying soil particle size and shape and drying rate are needed. Concerning the transport properties, simulations need to be done in three dimensions where the effect of connectivity on the diffusion coefficient is qualitatively similar, but quantitatively different. Similarly, the effect of sample size needs to be investigated. A challenge is the difficulty to image mucilage in soils while it still contains a significant amount of water. In Benard et al. (2019), the samples were scanned air-dry and only the final mucilage distribution was imaged. Scanning wet samples led to a destruction of the polymer network.

From a computational point of view, dynamic interactions between mucilage deformation, drying and solute transport within the network need to be implemented. Finally, the impact of microbial degradation and transformation of mucilage (see example 5) and the impact of mucilage on microbial activity need to be interconnected. Beside the complexity of such computational challenges, the experimental knowledge of such feedback between mucilage properties, transport processes and microbial activity is still in its infancy.

#### Example 3: Exudation by a growing root system

**Question** What is the effect of root architecture development and exudate properties on the 3D distribution of rhizodeposits in soil?

**Scale** mm-cm, days-weeks

#### Approach

We coupled the 3D root architecture model CPlantBox (Schnepf et al., 2018) with a rhizodeposition model to investigate the spatio-temporal distribution patterns of rhizodeposits in the soil as affected by root architecture development and rhizodeposit properties. To simulate the 3D dynamic pattern of rhizodeposits in the soil, we considered each growing root to be a moving point source or a moving line source. In the soil, rhizodeposits were subject to diffusion, sorption and decomposition. Microorganisms were not considered explicitly, but degradation of rhizodeposits was included in form of linear first order decay (Kirk et al., 1999).

Analytical solutions for moving point or line sources in an infinite domain have long been available (Carslaw and Jaeger, 1959). Thus, for any point in time or space, the analytical solution for rhizodeposit concentration around a growing root can be computed analytically. As the underlying partial differential equation is linear, the concentration due to multiple roots of a growing root system was then computed, using the superposition principle, as the sum of concentrations due to each root. The fact that the solution is based on analytical solutions does not mean a low computation time, however, because it involves the evaluation of integrals. Unless approximations are done that allow the analytical solution of those integrals, they have to be evaluated numerically as many times as the spatial and temporal resolution requires. In this example, the simulation time was set to 21 days, simulation outputs were generated every day. The size of the soil domain was 20×20×45 cm^3^, and we computed the concentration at every mm^3^. Simulations were performed for two rhizodeposits, mucilage and citrate and two different types of root systems, *Vicia faba* and *Zea mays*. All computations were performed on the Linux cluster of IBG-3 at Forschungszentrum Jülich, which facilitated parallelisation. Details can be found in Landl et al. (2021).

#### Parameterisation of the root architecture model

For the simulation of root architectures, CPlantBox requires a minimum of 13 parameters for each root type (i.e. for the primary and shoot-borne roots as well as for each order of laterals). These parameters describe the emergence, branching, elongation and death of the roots as well as their orientation in 3D space that may depend on different kinds of tropisms such as gravitropism, chemotropism or thigmotropism. Further optional parameters include the scaling functions for the root elongation, branch spacing and branching angle, e.g., in response to local soil conditions. In this example, root architecture parameters were derived from µCT images of *Vicia faba* and *Zea mays* roots (Gao et al., 2019). Six replicates of µCT scans were available for both *Vicia faba* and five for *Zea mays*. Each *Vicia faba* root system was scanned at 7, 11, 15 and 19 DAS, while each *Zea mays* plant was scanned at 11, 15 and 19 DAS. The roots were manually traced in the 3D virtual reality system available at the Supercomputing Center of Forschungszentrum Jülich, resulting in a data structure called root system markup language, (Lobet et al., 2015). It stores the 3D coordinates of nodes along the centre line of the root system and their connectivities, along with different properties for the edges (root segments) connecting two nodes, such as the root segment radius or age. The root segment ages were computed from linear interpolation between the measurement times using the respective elongation rates. Many model parameters can be directly computed from the RSML files, such as the mean and standard deviations of radii, apical and basal zone lengths, the intermodal distances or the branching angles. The remaining parameters such as the growth parameters and emergence times of basal roots were obtained by inverse estimation that minimized the difference between observed and simulated total root length.

#### Rhizodeposit parameters

Rates at which both plants release rhizodeposits (mucilage and citrate) were derived from literature (Rangel et al., 2010; Zickenrott et al., 2016). We assumed that mucilage was released at the root tips, while citrate exudation occurred at a length of 4 cm behind the root tips. The diffusion coefficients in water for mucilage and citrate were taken from Watt et al. (2006), the impedance factor from Olesen et al. (2001), the soil buffer power for citrate from Oburger et al. (2011) and the decomposition rate constants for mucilage and citrate from Kirk et al. (1999) and Nguyen et al. (2008).

Fig. 7 outlines the workflow of example 3. An important step in this example was the development of a model that can simultaneously simulate root architecture development and root exudation as well as the fate of root exudate within the soil. As in examples 1 and 2, data acquisition includes a plant growth experiment with a soil column. This time the µCT images were processed in such a way that roots are segmented and traced so that parameters for the root architecture model CPlantBox can be derived. From a second, independent experiment, rhizodeposition rates are determined for each individual rhizodeposit. Together with their respective transport and reaction rate parameters, the simulated outcome of this example is the 3D dynamic pattern of rhizodeposition concentration for different rhizodeposits and plant species.

**Fig. 7.**
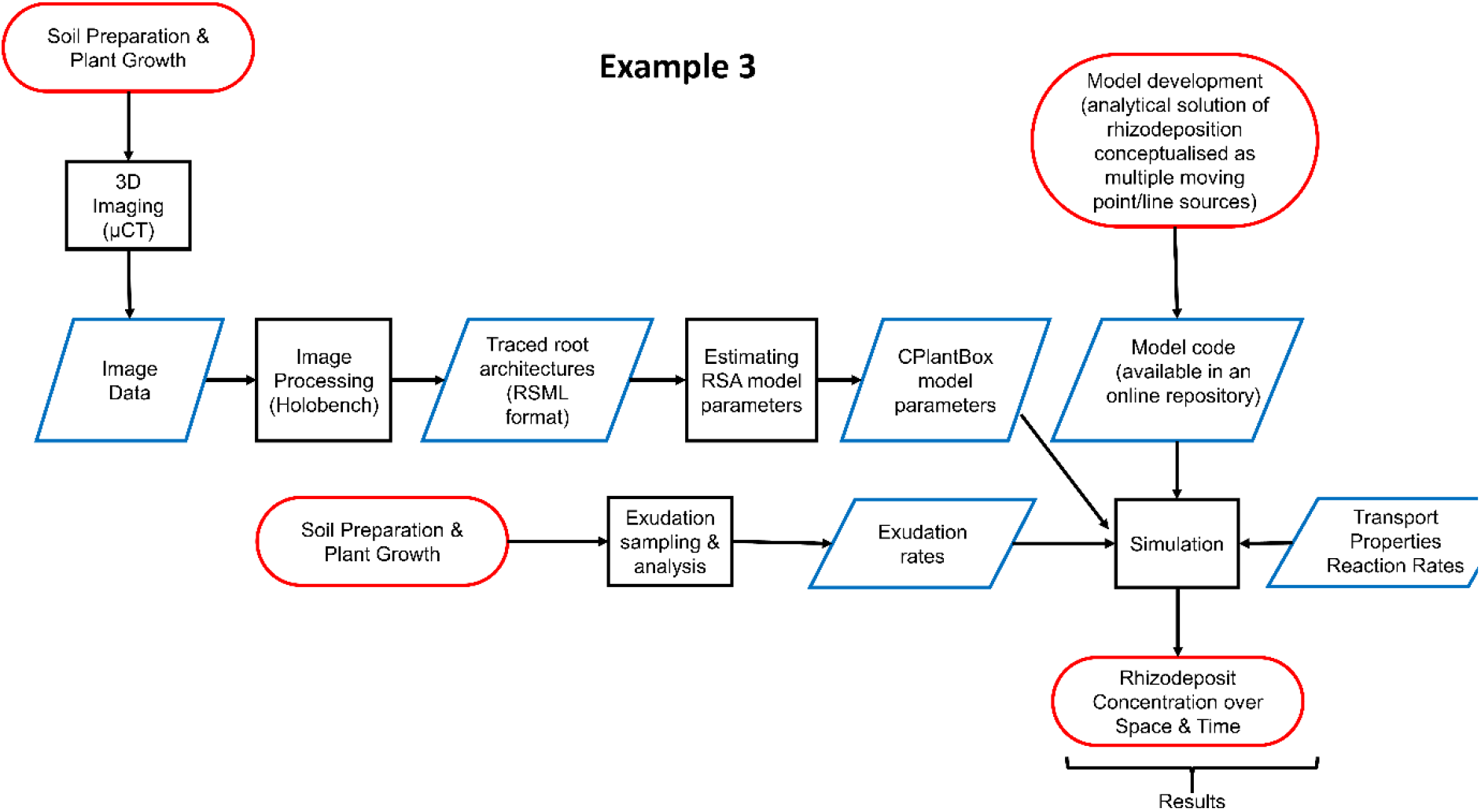
Flow chart outlining the approach of example 3

**Fig. 8.**
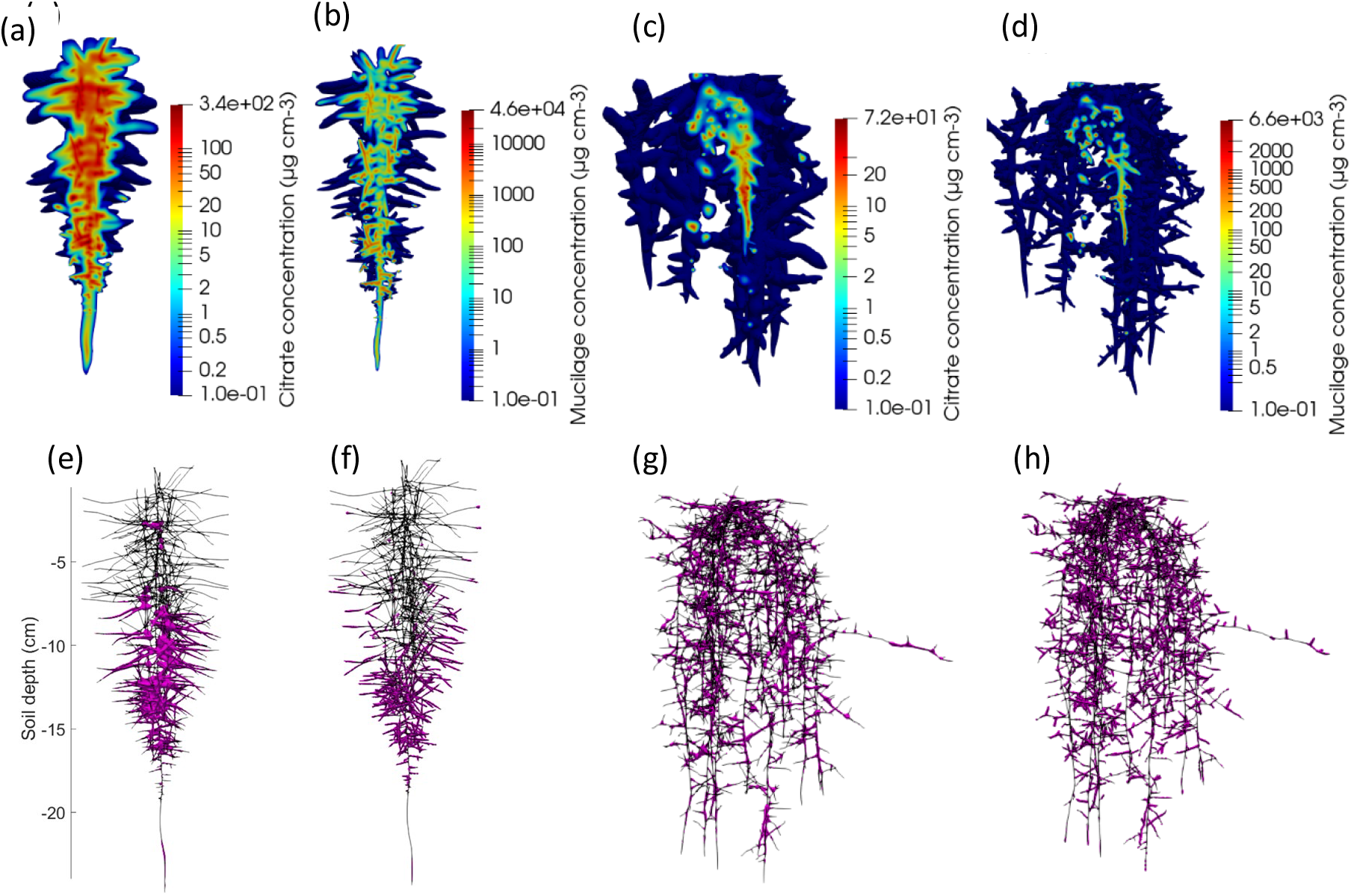
Vertical cut through the distribution of the rhizodeposit concentrations around 3-week old root systems (citrate around *Vicia faba* (a), mucilage around *Vicia faba* (b), citrate around *Zea mays* (c), mucilage around *Zea mays* (d)); note that the colours are in logarithmic scale. From Landl et al. (2021). Distribution of rhizodeposit hotspots (pink patches) of citrate and mucilage around a 21-day-old root system of *Vicia faba* (e, f) and *Zea mays* (g, h).

Fig. 8 shows a vertical cut through the distribution of citrate and mucilage concentrations around the 3-week old root systems of *Vicia faba* and *Zea mays*. It also shows corresponding distribution of rhizosphere hotspots, in which the rhizodeposit concentration is above a defined threshold. For mucilage, we chose a value at which the mucilage concentration is high enough to significantly alter soil hydraulic properties, 0.33 mg g^-1^ dry soil (Carminati et al., 2016). For citrate, we chose a concentration that is high enough to significantly mobilise phosphate (Lyu et al., 2016).

Our simulations showed that root branching allowed the rhizospheres of individual roots to overlap, resulting in a greater volume of rhizodeposit concentrations above a defined threshold value (i.e., the rhizodeposit hotspot volume). This was particularly important in the case of citrate, where the overlap of rhizodeposition zones accounted for more than half of the total rhizodeposit hotspot volumes. The rhizodeposit hotspot volume around the tap root system of *Vicia faba* was shown to be much larger than around the fibrous root system of *Zea mays*, which is due to differences in both root architecture and rhizodeposition rates.

#### Challenges and open questions

The combined effects of root architecture development and rhizodeposition in turn affect multiple other relevant processes such as soil carbon turnover and microbial activity or plant water and nutrient uptake. Modelling those interactions will enable us to assess the impact of rhizodeposition on these processes on the root system scale. Little information still exists on changes in the diurnal rhizodeposition and its dependence on light quality and quantity. New experimental data sets on the distribution of rhizodeposits in soil are now available, including zymography or co-registered Magnetic Resonance Imaging and Positron Emission Tomography (Koller et al., 2018). Integrating those data into our model, we will be able to increase our process understanding as well as gain additional information such as on the location, strength and temporal dynamics of rhizodeposition rates using inverse modelling.

#### Example 4: Phosphate uptake by a growing root architecture as affected by citrate exudation

**Question** How is overall phosphate solubility and consequently uptake affected by root exudates released from a growing root system? How well are diffusion, transport and reaction of nutrients in the rhizosphere of a growing root architecture understood?

**Scale:** mm-cm, days-weeks

#### Approach

The dynamic root architecture model CPlantBox (Leitner et al., 2010a; Schnepf et al., 2018) was coupled with a model of phosphate transport in soil and rhizosphere (i.e. the soil cylinder around each root segment) that takes into account competitive sorption of phosphate and citrate in soil. In particular, the sink term for phosphate uptake by roots from soil was developed in a way to recognize the dynamic development of the rhizosphere gradients around each individual root segment. For each root segment of the root system, a 1D radially symmetric rhizosphere model was solved in each time step and coupled to the macroscopic plant-scale model in a mass-conservative way. The model setup was a virtual representation of a rhizobox experiment with oilseed rape (*Brassica napus* L.) (see Schnepf et al., 2012 for details). Briefly, oilseed rape was grown in a rhizotron for 16 days and kept at an inclination of 49°. The simulations mimicked this experiment such that simulated root growth was confined inside the rhizotron geometry and had the same root length as observed, namely 734 cm. The root architectural parameters were obtained by inverse estimation such that the difference of observed and simulated root length densities was minimized. The water content was set constant at 0.3 cm^3^ cm^-3^. Root exudation was assumed to occur at the whole root length with an exudation rate of 3×10^−6^ µmol cm^-2^ s^-1^ (Kirk et al., 1999). A competitive Langmuir sorption isotherm for citrate and phosphate was used to simulate phosphate mobilisation due to citrate exudation. The sorption parameters, microbial decomposition rate constant for citrate, and Michaelis Menten P uptake parameters were derived from literature (see Tables 1 and 2 in Schnepf et al., 2012). Initially, the soil had a homogeneous initial P concentration and zero citrate concentration. At the boundaries of the rhizotron, no-flux boundary conditions were prescribed, so that the only changes in citrate and phosphate concentration in the rhizotron occurred through the root activities, citrate exudation and P uptake.

**Table 1.**
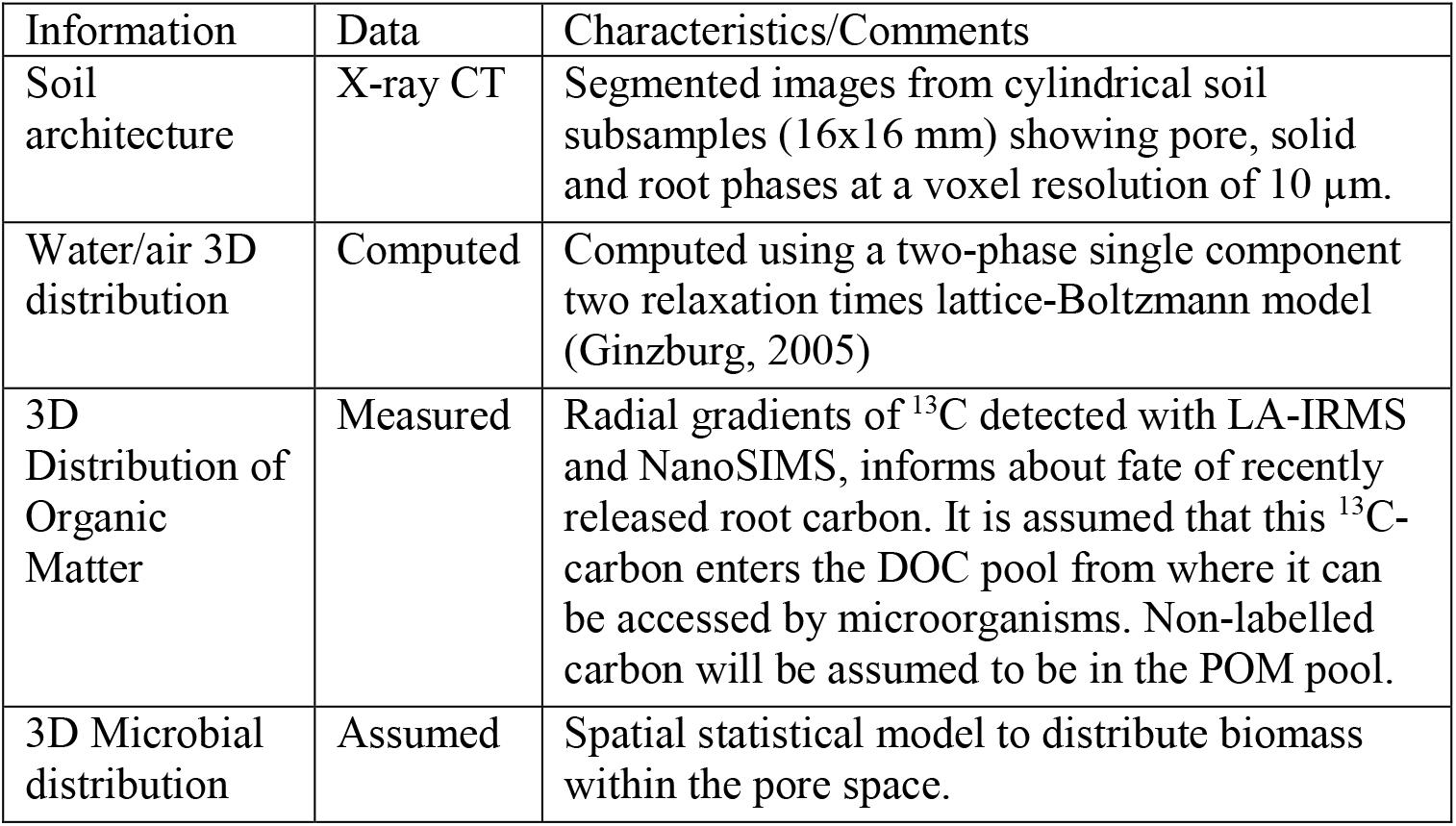
Information required to initialise a modified Portell et al. 2018 model including root exudates.

| Information | Data | Characteristics/Comments |
| --- | --- | --- |
| Soil architecture | X-ray CT | Segmented images from cylindrical soil subsamples (16x16 mm) showing pore, solid and root phases at a voxel resolution of 10 $\mu\text{m}$. |
| Water/air 3D distribution | Computed | Computed using a two-phase single component two relaxation times lattice-Boltzmann model (Ginzburg, 2005) |
| 3D Distribution of Organic Matter | Measured | Radial gradients of $^{13}\text{C}$ detected with LA-IRMS and NanoSIMS, informs about fate of recently released root carbon. It is assumed that this $^{13}\text{C}$-carbon enters the DOC pool from where it can be accessed by microorganisms. Non-labelled carbon will be assumed to be in the POM pool. |
| 3D Microbial distribution | Assumed | Spatial statistical model to distribute biomass within the pore space. |

Fig. 9 outlines the workflow of example 4. Model development resulted in a model that could simultaneously simulate root growth, root exudation and its effect on nutrient availability and uptake. Data acquisition involved a plant growth experiment where roots were washed from the soil, scanned, and root length determined with WinRhizo. This data was used to inversely estimate root architectural parameters for the CPlantBox model. Together with transport properties and reaction rate parameters, the simulation results in the 3D dynamic pattern of phosphate and citrate concentration as well as cumulative nutrient uptake and exudation.

**Fig. 9.**
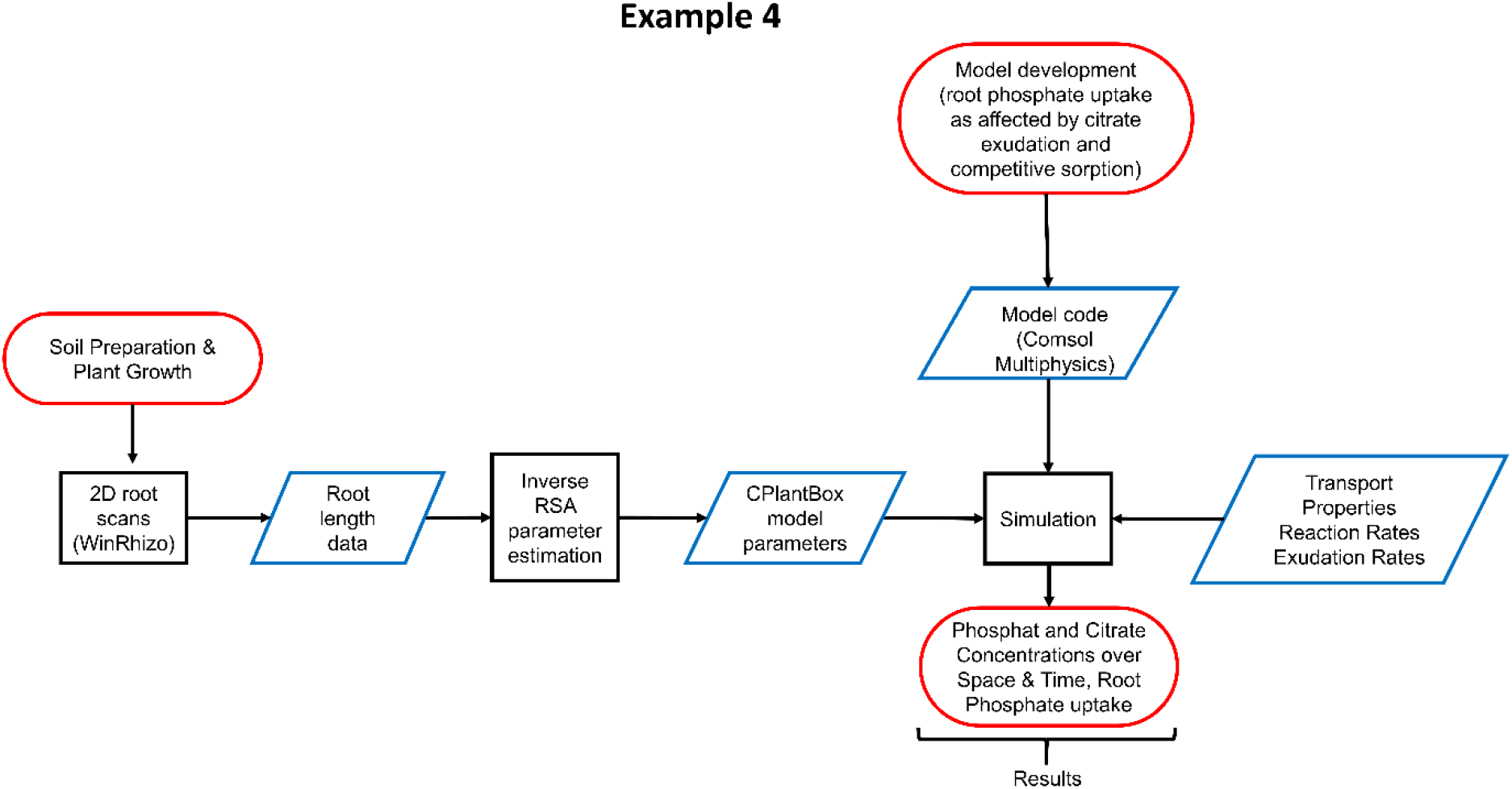
Flow chart outlining the approach of example 4

## Results

This model quantifies how two traits, root system growth and exudation, affect plant phosphate uptake. Fig. 10 shows the age distribution in the simulated root architecture of roots grown inside a rhizobox, as well as the total phosphate and citrate concentrations in soil as affected by root exudation and phosphate uptake. We can see that the highest citrate concentration and phosphate depletion occurs in a region where the root system is oldest. In this example, performed for a soil with medium sorption capacity, the model showed that cumulative phosphate uptake was more than doubled through root exudation of citrate (see Schnepf et al., 2012). This value would vary for soils with different sorption behaviours. The example shown here is based on the assumption that root exudation occurs over the whole root length, so that the citrate concentration would be higher around older roots. This is different if the root exudation occurs only near the root tips as we have seen in example 3, as then concentrations are highest near the root tips. In both cases, overlapping accumulation zones would further induce hotspot formation.

**Fig. 10.**
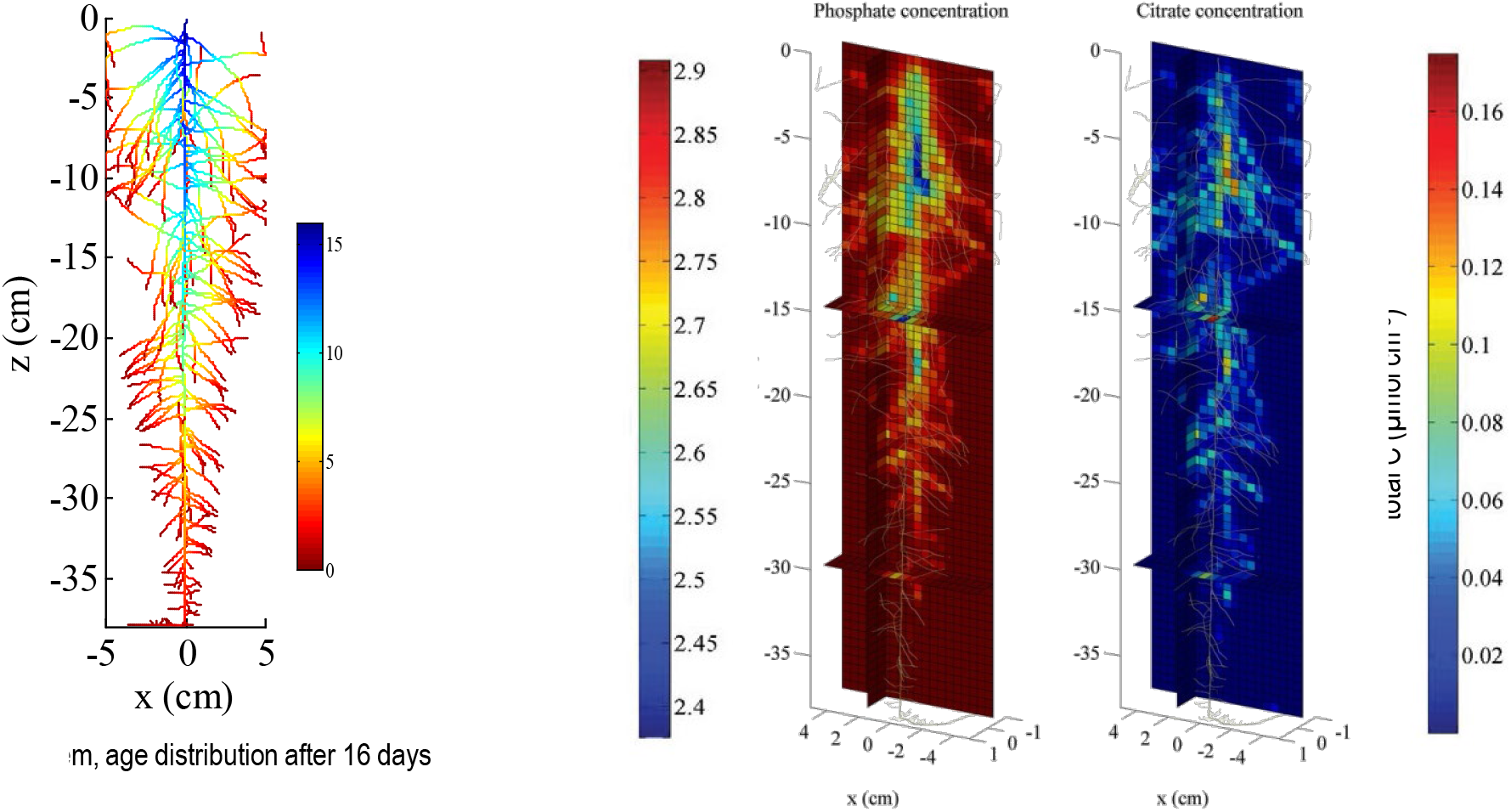
Left: Simulated root architecture of oilseed rape, colours denote root segment age. Right: Total phosphate and citrate concentrations in soil after 16 days of simulation when root exudation occurs over the whole length of the root axes. From Schnepf et al. (2012).

### Challenges and open questions

Functional-structural models of root water and nutrient uptake may help to understand what would be the optimal coordination between root growth, rhizodeposition of different substances to obtain optimal water and nutrient uptake by the root system. The simulation results presented here made assumptions about the location and dynamics of exudate release along the root axes. Now available experimental data will help to a more accurate parameterisation. Another alternative is the coupling to whole-plant structural functional models in which the flow of carbon from photosynthesis to the different plant organs and release into the soil is explicitly modelled. Those model results may be used as input for the model presented in example 3.

The effective diffusion coefficient used to simulate nutrient diffusion in soil was computed using the standard Millington-Quirk model (Millington and Quirk, 1961). A current challenge is to derive an effective diffusion coefficient as a function of radial distance from the root surface using mathematical homogenisation based on µCT-derived soil structure and mucilage information as shown in example 2.

Another challenge is to compare the simulated nutrient gradients to the gradients found through chemical imaging methods at different positions within the root system and to evaluate whether the model can reproduce those observed gradients. If this is established, then observed element distributions could serve as input for inverse simulations for nutrient uptake and transport parameters.

### Example 5: Microbial activity based on C-distributions: using modelling to integrate rhizosphere processes

**Question** Can we identify the key drivers of microbial activity and C distribution in the rhizosphere considering a holistic approach?

**Scale** from nm to cm; seconds to days/weeks.

### Approach

A modelling framework is proposed that can predict microbial dynamics and processes resulting from interactions in the soil surrounding roots.

We conjecture that the rhizosphere as we measure it emerges from interactions occurring at the pore-scale around roots. The complexity of the rhizosphere demonstrated by the examples in previous chapters may appear overwhelming at first sight, but we propose that it is possible to fully embrace this by a new generation of mechanistic, spatially explicit pore-scale models, building on recent advances in this area. For instance, using a mechanistic pore-scale fungal model, it has been shown that fungal growth in soil is non-linear and that, for a given volume and a given nutrient content, it depends on the micrometre scale distribution of nutrients and microbes (Falconer et al., 2015a). These micro-scale heterogeneities explain nonlinearities in the temporal evolution of fungal biomass, carbon degradation and CO_2_ flux observed experimentally. Focusing on soil bacterial dynamics, recently, it has been demonstrated how heterogeneity at the micro-scale can favour poor bacterial competitors, suggesting that pore geometry is a driver of soil biodiversity (Portell et al., 2018). These results indicate that mechanistic (first-principle) models are essential for upscaling microbial driven processes and that this approach can help to bridge the gap between pore-scale and the continuum-scale description of the system (Vetterlein et al., 2020). In this section, we propose the modification of an existing pore-scale microbial soil model as a first step to integrate rhizosphere experimental data and highlight the benefits and limitations of this approach and the path ahead for further work.

#### Proposed modelling framework for rhizosphere behaviour: an individual based approach

Our starting point can be a spatially explicit, pore-scale model accounting for the activity of soil bacteria (Portell et al., 2018). Briefly, Portell et al. (2018) assumes a cubic lattice describing explicitly the soil architecture where particulate organic matter (POM) and microorganisms are located. POM hydrolyses over time creates dissolved organic carbon (DOC) that is released to the water phase where it becomes available for bacterial growth or diffuses away to more distant areas as determined by a lattice-Boltzmann model component. Bacterial position and physiology are controlled by an individual-based approach accounting for single bacteria that divide when the cell attain a critical mass.

In the rhizosphere, organic matter distribution can be seen as the result of the distribution of two main components that can already be mapped to organic matter pools considered by Portell et al. (2018). The dominating fraction is a large and dynamic C fraction that is continuously exuded to the soil from the region between root tip and root hair zone and that can be mapped to the DOC pool of the model. The distribution of “dynamic C” (i.e., root-exuded C) follows a gradient into the soil and can be traced using stable C isotope labelling. Labelled C exuded by the plant root can be followed and quantified on the millimetre-scale using an adequate sampling technique and can be visualized in 2D at the micro-scale when using resin embedded root-soil sections and nano-scale secondary ion mass spectrometry (NanoSIMS) (Vidal et al. 2018) or laser ablation-isotope ratio mass spectrometry (LA-IRMS) (Rodionov et al., 2019). This information can be used following two different approaches. In the first approach, a boundary condition is placed on the root surface of the living roots obtained from CT images (Ruiz et al., 2020) to account for the exudation of carbon from the root. Following this approach, the gradients measured will be used to calibrate the exudation parameter of the model. In a second approach when experimental information is available, we assume a spatially static gradient of exuded C.

The second fraction of C in the rhizosphere is a rather static, discontinuous fraction made of dead plant residues and microbial necromass and mineral-associated organic matter (MAOM) (e.g., Liang et al., 2019) that can be mapped to the POM considered by Portell et al. (2018). This fraction, which we can expect to be located more irregularly in areas with decreased accessibility by the microorganisms (Rodionov et al., 2019; Totsche et al., 2018) can also be estimated using NanoSIMS and LA-IRMS.

In bulk soil, bacteria invest energy to produce enzymes that hydrolyse POM over time, thereby releasing DOC to the water phase where it becomes available for bacterial growth or diffuses away to more distant areas (as dictated by the lattice-Boltzmann component of the model). It must be noted that the majority of soil microorganisms has limited access to available carbon and resides in a dormant state most of the time. In the immediate vicinity of the growing root tip however, the release of DOC via exudation transiently lifts the carbon limitation of microbial growth in soil and favours fast-growing copiotrophic taxa (Bonkowski et al., 2021; Rüger et al., 2021). Modelling these processes requires accounting explicitly for trade-offs associated with the release of enzymes by microorganism, a function currently assumed ubiquitous in the model, as well as the diffusion of released enzymes through the water phase from where they can reach and control hydrolysis of distant volumes, where it might initiate a priming effect on POM.

Experimentally it has been observed that the microbial growth rate is a function of the availability of C-containing exudates, whose diffusion rates differ among molecule classes (Jones et al., 2004), therefore showing effects dependent on the distance and location of the root surface in relation to the point in space considered. As a first approximation, models could be based on a uniform source of DOC. At suboptimal DOC levels, carbon has been shown to be respired without any microbial growth (Anderson and Domsch, 1985), while higher DOC levels, especially in the close vicinity of roots stimulate microbial growth. High DOC levels may also favour microbial enzyme production that subsequently leads to enhanced hydrolysis of carbon and nutrients from POM, a self-enhancing positive feedback mechanism on microbial growth, known as priming effect (Kumar et al., 2016; Mo et al., 2021). Assuming the individual-based model approach, this can be simulated with the implementation of appropriate rules ensuring that the growth and enzyme production is only possible when microbial maintenance requirements are covered. In addition, current root growth models suggest that microbial attachment to roots is a key strategy to gain maximum access to rhizodeposition (Dupuy and Silk, 2016). Therefore, next to the growth rate, future models must consider microbial motility. Microbial motility is disregarded in the work of Portell et al. (2018), although the model structure is designed to allow the implementation of such modification.

Recently, Dupuy and Silk (2016) studied the colonisation of root surfaces resulting from the growth of root tips in bulk soil. These authors modelled root growing through a homogeneous (continuous) soil fraction and used a population-based approach to account for microbial growth and colonization dynamics. Dupuy and Silk (2016) found that the root elongation rate was a key trait for successful establishment of bacteria on the root surface. This suggest that in addition to microbial growth and motility, future models should consider root growth in order to reflect more realistic colonization dynamics. When structural information of the solid, water and air phases of the soil are taken into account, as we are suggesting here, the introduction of growing roots in these models is challenging. In certain situations, time-lapse imaging could be used to follow root growth and its modifications of soil structure in order to implement this into complementary modelling scenarios. To our knowledge, this has not been achieved to date. Instead, root (or root hair) growth is tackled by introducing time dependent boundary conditions. In this approach, a common technique is to use a fully grown geometry and to activate the appropriate boundary conditions according to a (measured) growth rate (e.g. McKay Fletcher et al., 2020). Given the impossibility of following root growth in the field, we advocate the use of an equilibrium approach and static roots as a first approximation. Given the complexity of the rhizosphere, this is also advisable. We need to understand rhizosphere mechanisms around static (or slowly growing roots) before embracing the full complexity of the system.

### Simulation scenarios and model initialisation

Structural information supplemented with existing knowledge will describe the physico-chemical state of the soil microhabitats around roots and be used as initial condition to conduct spatially explicit simulations with the pore-scale model Fig. 11a).

**Fig. 11.**
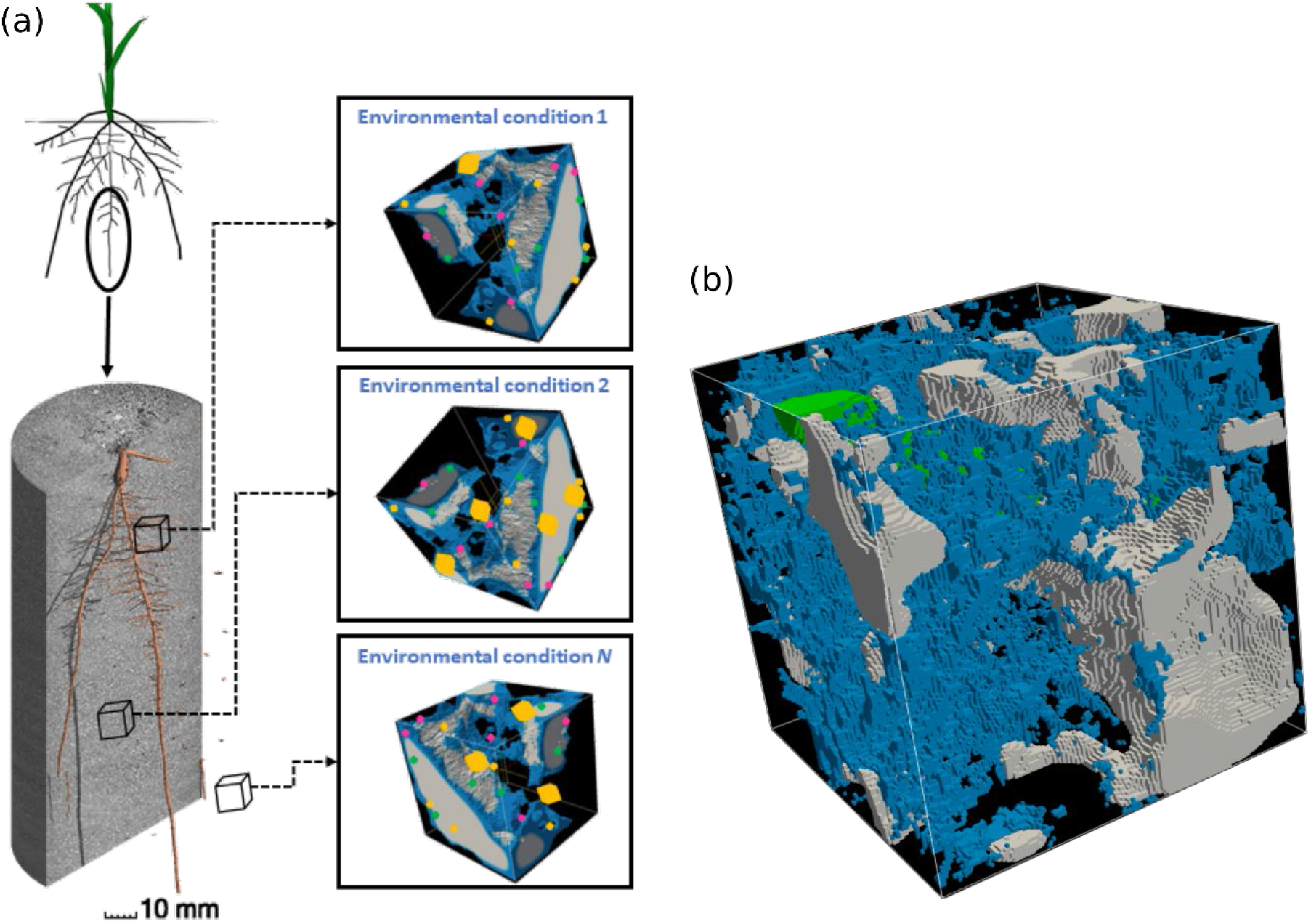
Scheme of the simulation scenarios (a) and detail of a 128^3^ sub-domain showing segmented roots (green) and the water (cyan) and air (grey) phases computed using the lattice-Boltzmann model described by Genty and Pot (2013) assuming a water saturation level of 50% (b).

The essential information required to initialise the model is summarised in Table 1. A number of steps are required to set the physical environment surrounding roots. The soil architecture of the rhizosphere, including root positions, will be obtained with segmented X-ray CT images. Smaller sub-domains (e.g., 128^3^ voxel size) will be obtained from different distances from the root surface (Fig. 11b). The first challenge is to simulate the distribution of water and air within the pore space of the subdomains as described in the example 2 above.

Root derived C has been shown to be distributed at least 100 µm distant from the root plane (Rodionov et al., 2019), but more likely extends in the mm range (Jones et al., 2004). Our first working assumption will be to assimilate 13-C labelled C to diffusable dissolved organic matter (DOC, i.e., readily available C for microorganisms) and all non-labelled C within a 100-1000 µm-rhizosphere-range as particulate organic matter (POM, that can be mineralized under the action of microbial enzymes).

The initial physico-chemical environment described so far will be the set up where the microorganisms evolve as controlled by the individual-based bacterial model. The 3D distribution of microorganisms has been obtained in bare soil (Eickhorst and Tippkötter, 2008; Juyal et al., 2019; Nunan et al., 2001), but is largely unknown to date for the rhizosphere. To account for this experimental lack, our approach will use a preliminary simulation initialised using the bacterial numbers and distributions found in the bare soils. Since the microbial biomass and activity is modulated by microbial traits and the carbon available in the media, the microbial numbers reached in the preliminary simulation will depend on the biochemical conditions of the media. Spatial distributions of bacteria in soil thin sections have been approached using a 2D spatial statistical model (Raynaud and Nunan, 2014) that can be expanded to a 3D soil structure. Bacterial biomass and respiration will be monitored in all spatial grid elements and plotted as a function of the distance to the root plane. The microbial model can be calibrated using quantitative measures of microbial respiration (i.e., mineralization) of different quantities of root exudates on bare soil such as the measurements shown in Fig. 12.

**Fig. 12.**
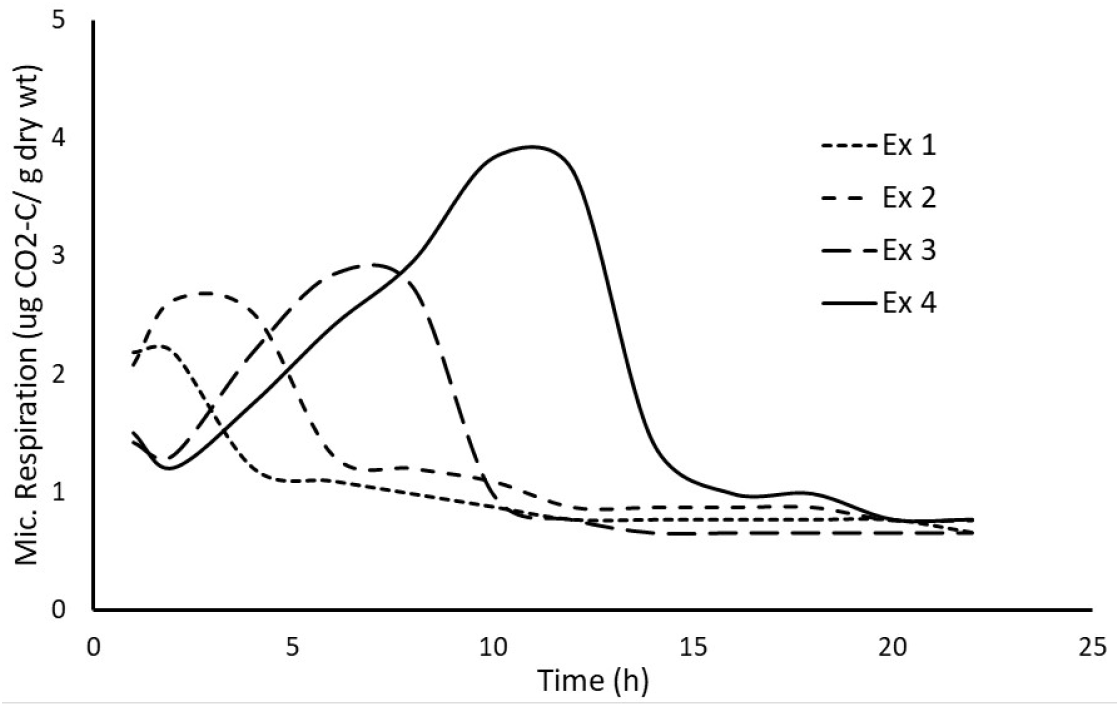
Microbial respiration in soil amended with increasing quantities of exudates (Ex 1-Ex 4). Note that the lowest exudate quantities (Ex1, Ex 2) only led to a short-term stimulation of microbial respiration (but not growth), while respiration curves with higher concentrations (Ex 3, Ex 4) show typical microbial growth dynamics.

Model outputs will be qualitatively validated using microbial biomass and respiration measured at 3 mm intervals from the rhizosphere plane (Alphei et al., 1996).

A flow chart outlining steps from experimentation to the generation of model outputs can be found in Fig. 13. It combines the workflow of example 2 (resulting in water distribution in 3D pore space) with carbon distribution as measured from resin-impregnated soil samples derived from the same experimental setup and the initial bacterial spatial distribution as simulated from a bacterial spatial statistical model. Together with the appropriate boundary conditions, transport properties and microbial behaviours and properties the simulation results in a 3D dynamic system where biotic and abiotic components interact. Results can be aggregated to provide continuum-scale properties such as bacterial mass or respiration per soil control element as a function of time.

**Fig. 13.**
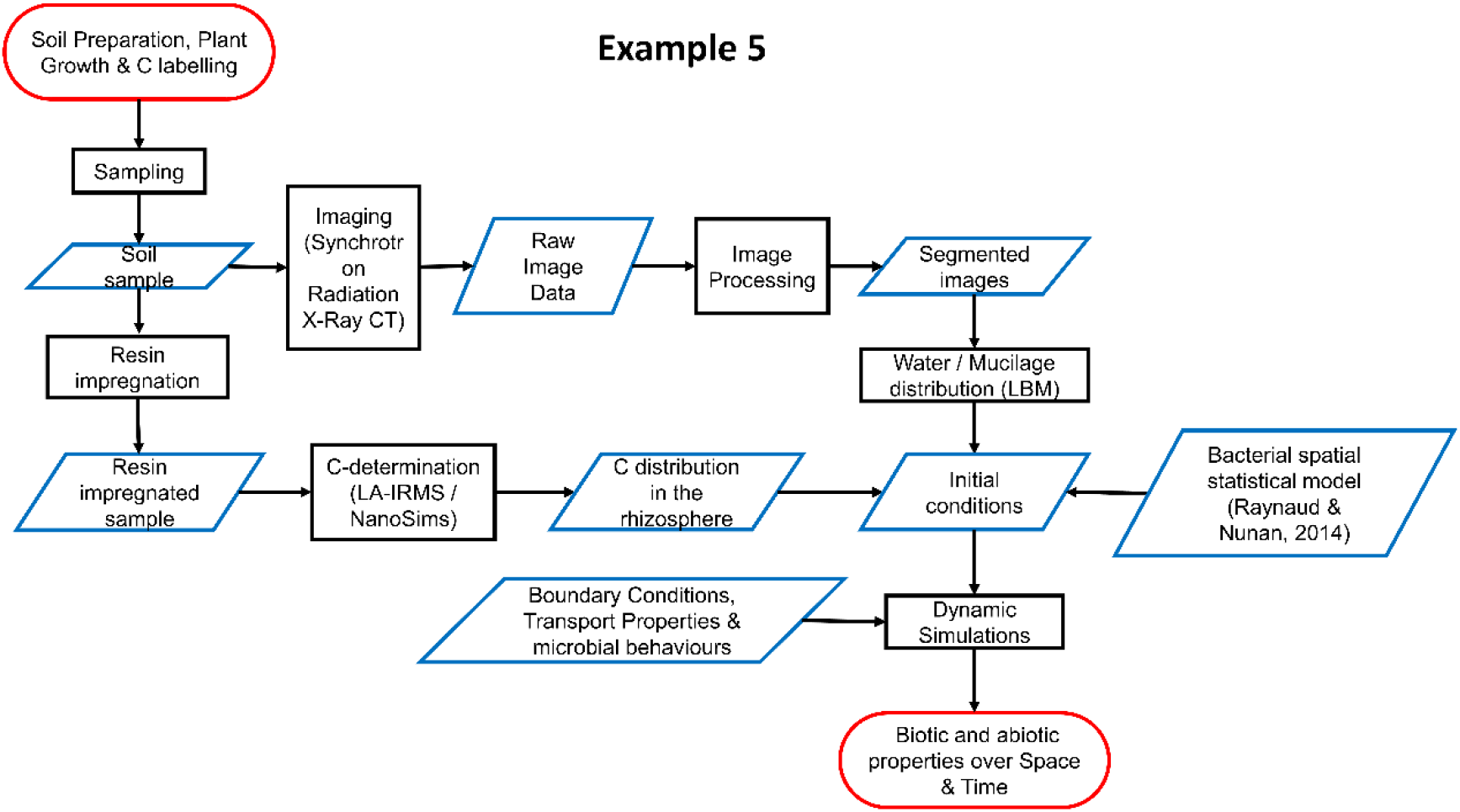
Diagrammatic flow chart of example 5. In the figure, CT stands for Computed Tomography, LBM stands for lattice-Boltzmann model, µIbM stands for microbial individual-based model, LA-IRMS stands for laser ablation-isotope ratio monitoring, and NanoSIMS stands for nanoscale secondary ion mass spectrometry.

## Results

The approach highlighted here is a first step towards the development of mechanistic models accounting for C distribution, and C transformation by microbial activity around roots. Assuming an equilibrium approach and one static root allows us to integrate the various experimental data collected and use them to predict the shape and distribution of the microbial activity in the soil surrounding roots and, more importantly, allows identification of the abiotic components or gradients measured driving the rhizosphere shape and extend. The approach is also a steppingstone towards the development of the much needed, fully fletched pore-scale rhizosphere model (see challenges and open questions section below).

The approach discussed in this section offers spatially explicit information of the bacterial biomass, the carbon decomposition, and the evolution of bacterial respiration over time. This allows estimating microbial activity namely biomass, respiration and spread rate as a function of the distance from the root surface. The use of an individual-based approach allows also to tackle fundamental and applied question related to the maintenance and development of microbial diversity in the rhizosphere.

### Challenges and open questions

Rhizosphere complexity requires accounting for more reactive chemical species. For instance, recent publications highlight the important role of volatile carbon signals that expand the rhizosphere in the cm range (de la Porte et al., 2020). In addition to dissolved organic carbon (e.g., glucose), inclusion of nitrogen containing organic compounds (peptides, amino acids), CO2, O2, and signalling with phytochemicals such as lipo-chitooligosacharides (Venturi and Keel, 2016) would allow to study a number of important mechanisms underlying microbial colonisation and establishment in the rhizosphere.

Further differentiation of DOM into specific root exudates, microbial biomass, microbial necromass and POM (Angst et al., 2016; Baumert et al., 2021) can be envisaged and tackled with the same modelling approach for an increased accuracy of the model outputs. This approach may even allow the characterization of priming effects in different aggregate size classes in the rhizosphere based on microbial stoichiometry (Mo et al., 2021; Wang et al., 2020). Molecular marker analyses including a suite of exudate-, bacteria- and fungi-specific substances has high potential to shed more light on rhizosphere gradients and hot spots of OM enrichment (time-integrated signals/stable on medium-term in contrast to microbiological parameters) (Baumert et al., 2021). As a first step ^13^C labelled carbon fraction which can be imaged in 2D with NanoSims or LA-IRMS is used to represent the soluble root derived carbon pool (exudates) and complement the C initialisation described and/or to assist validation of experimental results, especially in view of stoichiometric constraints for plant and microbial processes (Clode et al., 2009; Gorka et al., 2019; Vidal et al., 2018).

This first adaptation of existing spatial-explicit mechanistic models for the rhizosphere follows an equilibrium approach and one static root. A fully fletched pore-scale rhizosphere model requires the explicit representation of a dynamic root capable of simulating uptake and release of water and elements from (to) the soil. In addition to a static water flow scheme, water dynamic can be also taken into account using LBM approaches in both saturated and unsaturated porous media (Ginzburg, 2008; Zhang et al., 2016).

Another fundamental issue comes from the huge, and largely unknown, microbial variability existing in soil. We advocate for the use of individual-based modelling of soil microorganisms in conjunction with a trait-based approach coupling bacterial (Portell et al., 2018) and fungal (Falconer et al., 2015a) models. Bacteria and fungi act at different scales and prefer different soil phases (i.e., water against air filled porosity), which suggest the need of this first division. Although the whole soil biodiversity is intractable using a species-centric approach, a trait-based approach can be used to identify trade-offs in microbial traits and strategies that lead to an improved rhizosphere colonisation. Such information has practical implications by extrapolation to specific species. For bacteria, we suggest accounting for the following minimum set of individual traits: lag phase length, specific growth rate, ability to grow anaerobically, mortality rate (by predators), motility, antibiotic production, root attachment behaviour, and C specificity (DOC, Mucilage, and plant signalling molecule –e.g., malic acid like). The consideration at the single-cell level of these traits would provide insights on beneficial traits and trait combinations for rhizosphere colonisation and establishment on root surfaces. For fungi, physiological trait combinations representative of functional groups proposed for arbuscular mycorrhizal fungi (Chagnon et al., 2013): competitors, stress tolerators, and ruderals, can be adopted. The first challenge is the identification of model parameters describing these experimentally unknown behaviours. Relevant questions are: what are the C costs vs. nutrient and/or water gains associated with the establishment and maintenance of mycorrhiza by roots? The adoption of an approach describing individual fungi with different properties and behaviours (Falconer et al., 2015b) will also allow to model interactions between fungal species such as the competition among symbionts (benefiting from root C) and necrotrophic pathogens that gain access to plant C only after killing parts of the root (Sarkar et al., 2019).

### Synthesis and Outlook

#### Detailed small-scale information to predict large-scale emerging behaviour

We have presented a series of examples illustrating the effect of rhizosphere scale properties affecting transport across the root-soil interface and microbial activity. We have chosen the example of exudation because of its relevance for nutrient uptake and microbial activity. We have illustrated how root hairs, at a given exudation rate, can increase the total exudation as well as the spatial extent of rhizodeposits into the soil. We have then discussed the mechanisms by which the polymers present in mucilage increase the viscosity of the soil solution and thus alter the spatial configuration of the liquid phase. Specifically, the high viscosity mucilage prevents the break-up of liquid bridges between soil particles and maintains the diffuse pathway for solute across the rhizosphere. The effect of root hairs and mucilage on solute transport across the rhizosphere could then be implicitly described with effective diffusion coefficients, as we have shown for the case of mucilage. Such effective diffusion coefficients can be implemented in root architecture models, where the effect of local rhizosphere properties (defined at the scale of root segments) can be investigated at the root system case. We have presented two examples, one in which we have calculated the spatial extent of different rhizodeposits across the rhizosphere and along the root system, and one in which we simulated the effect of root exudation on root nutrient uptake. Finally, we have presented a pore-scale example of a model of microbial activity, which takes explicitly into account the spatial distribution of soil particles, water, carbon and microorganisms. Such a model allows accounting for emergent microbial driven processes emerging from large number of biotic and abiotic interactions occurring at the microscale, informing models using coarser spatial resolutions. At the same time, this model would benefit from the other examples described as it depends on the connectivity of the liquid and gas phases as well as on the spatial distribution of rhizodeposits.

The examples described in this contribution show how small-scale rhizosphere traits (such as root hairs and mucilage) impact the properties (e.g. diffusion) of the rhizosphere at the root segment scale, and how such properties could be implemented into a root architecture model to investigate root system scale output (e.g. total uptake of nutrients). This is an example of a bottom-up approach in which small-scale processes taking place at the pore scale are included into a macroscopic model defined at the continuous scale. Such upscaling is done by means of defining effective properties, as we have illustrated for mucilage, and as can be done for root hairs using the homogenization method (Leitner et al., 2010b; Zygalakis et al., 2011). Effective properties depend on the spatial arrangement of soil particles and of the liquid phase, which could be imaged *in situ*, for instance using X-ray CT. The price to reach a spatial resolution needed to resolve the pore space is that the field of view of such images is limited and might be below the REV (representative elementary volume) needed to define properties (such as diffusion) at the continuous scale. However, we argue that the spatial extent of some rhizosphere properties, for instance the spreading of mucilage in soils is smaller than the REV – i.e. it might cover only a few layers of soil particles. Therefore, it is allowed to define effective properties for such processes while having in mind that such transport properties cannot be used for longer distance transport. We illustrate this concept with an example relate to water flow. Consider a 2D heterogeneous medium composed of elements with varying conductivity distributed randomly and with the conductivity lognormally distributed. If the medium is large enough compared to the size of its elements, the effective conductivity of the discrete medium is the geometric mean of the conductivities of the elements. However, if the domain is thin, or the flow process takes place only in a thin section of the domain, then the effective conductivity that should be taken converges to the arithmetic mean of the conductivities (von Jeetze et al., 2020). For the rhizosphere, different effective diffusion properties should than be defined depending on the spatial extent of the process. To our knowledge, this problematic has not been addressed.

Beside the complexity of defining and determining effective properties, the advantage of such properties is that they can then be implemented in larger scale continuous models in which root architecture and root growth can be explicitly modelled. This bottom-up approach allows testing scenarios and qualitative behaviours. For instance, it can be used to investigate in what conditions and for what processes (e.g., water uptake, P uptake, etc.) small-scale processes that affect the local transport properties in the rhizosphere are relevant on the plant scale. Such analysis can be done only at the plant scale where the appropriate boundary conditions and vertical gradients in soil variables are explicitly simulated. A problem with posing this question is that the meaning of ‘relevant’ needs to be defined. An at-hand definition would be that using bulk soil properties instead of considering the rhizosphere properties for predicting processes at the plant level might lead to incorrect estimates. This means that plant system scale processes are sensitive to rhizosphere properties. These sensitivities may also be dependent on the environmental conditions. For instance, in wet soils, the impact of mucilage on water uptake is not important whereas it is in dry soils. If some roots of the root system still have access to wet soil, root water uptake by the entire root system will hardly be affected by the impact of the mucilage around roots in the drier soil layers. When the sensitivity question is answered positively, the next question is whether the same prediction of processes at the entire plant system scale could be obtained by using different bulk soil properties or using different root system properties. If the answer to this question is yes, then an improper representation of rhizosphere processes or properties could be simply ‘compensated’ by adjusting root or soil properties. The question translates to whether sensitivities of the behaviour at the plant scale to rhizosphere properties, root system properties, and soil properties are correlated or not. It implies that when processes are observed at the plant system scale, rhizosphere properties cannot be derived (e.g., by inverse modelling) without knowing the other correlated properties. In order to unravel the ‘relevance’ of rhizosphere properties and processes for plant system scale processes, sensitivity analyses with multiscale models could be carried out and used to identify interaction effects (i.e., when the change in the larger scale process is larger than the sum of the changes due to changes in rhizosphere properties, bulk soil properties and root properties). Conditions when such interaction effects occur could then be used to infer rhizosphere properties from the observed emergent behaviour at the larger scale.

However, upscaled simulations require not only the effective properties in the rhizosphere at a given point in space and time, but also information on how these properties evolve over time and along the root system. This requires then a top-down approach and a general cross talk of models at different scales: the large scale to define the boundary conditions and state variables for the small-scale process, and the small scale to estimate the effective properties for the large-scale model. The behaviour of the system emerges from the interactions between the scales.

#### The importance of interactions between different processes

In addition to the interactions between scales, the emerging rhizosphere behaviour is also the result of the interactions between different simultaneously occurring processes. Here we have focussed on the example of rhizodeposition, and its interaction with root growth, soil water flow, nutrient uptake, and microbial growth and respiration. However, these are still limited examples that could be extended. More generally, nutrient uptake affects the plant status and most likely the quality and quantity of exudation, resulting in a feedback loop. Root water uptake takes also part to such feedback, due to convective fluxes affecting solute distribution and to the gradients in soil water potential and soil moisture around the roots, which in turn affect diffusion. De Bauw et al. (2020) illustrated that interactions between water uptake and nutrient uptake at the plant level scale could be linked to a combination of rhizosphere transport and root system scale water uptake. Understanding and properly implementing into models interactions between these processes, and their feedback with plant growth is a key to predict the emergent effect of rhizosphere properties on the plant scale. While there is an understanding on the effects of certain drivers such as drought on individual rhizosphere processes, the effects of these drivers on multiple, simultaneously occurring processes are still poorly understood. This requires the extension of existing models to a larger number of rhizosphere components.

#### How can measurements be used in modelling? State of the art and challenges

If the question could be framed reversely, it would be simple to answer, at least for the scientists conducting experiments in the real world. The expectation is that models: (i) integrate different findings, (ii) enlarge in ‘in silico’ experiments the number of drivers to be tested or at least increase the parameter range within one driver, (iii) scale up from local, or small scale observations to the whole system, or the reverse if only bulk information is available and local information is of interest, (iv) provide high temporal resolution even if measurements could only be provided for a limited number of time steps, and finally (v) test if hypothesised co-occurring mechanisms/processes could bring about spatial or temporal patterns observed, in particular if they are non-linear in nature.

In the frame of a current priority program focusing on rhizosphere spatiotemporal organization (PP 2089, https://www.ufz.de/spp-rhizosphere/) it was hypothesized that spatiotemporal patterns in the rhizosphere are formed as a result of local interactions and numerous feed-back loops (self-organisation processes) and that these patterns result in emerging system properties for the soil-plant system. Scientist were encouraged to work with the same experimental platforms with a very reduced number of drivers (two textures, two genotypes, four growth stages, few compartments) in order to provide as many input parameters for the different models operating at different scales. Can this be a successful strategy? The following sub-sections will tackle this question.

##### Parameters for bottom-up as well as top-down modelling approaches

The focus on joint experimental platforms with a few drivers only, provides for the same system information at a range of scales. If we take carbon flow as an example, the experimental data range from changes in standing above and below ground biomass in the field during the whole growth season, to root system exudation rates and their change with growth stage, to spatial distribution of recently assimilated carbon in the vicinity of roots at the µm scale. Thus, data from different scales can be used as input parameters and for validation, respectively. For all these scales, spatially resolved, quantitative and qualitative information is available; however; the level of detail changes with the scale and the units cannot always be converted directly into each other.

##### Experimenter can never provide all the required data – bottom-up

In describing the rhizosphere, we are forced into high-resolution data acquisition because some of the players are very small and some of the gradients are very steep. This is complicated by the fact that the closer we look, the more complex things can become. While we move from the continuum scale to the pore scale for soil, we must move from the single root scale down to the tissue scale for roots, and seemingly well-established knowledge, such as smooth gradients extending from the root surface into the root (what is happening in the plant), are challenged by questions such as where along the radial gradient is uptake or release actually taking place within the root, where along the root do we have to measure, which is the tissue concentration relevant for calculating ^13^C diffusion into the soil – ^13^C concentration of tangential walls of the endodermis or rather the mean ^13^C concentration of grind up root tissue from the whole root system? What adds to the experimenter’s dilemma is that the higher the resolution, the more cumbersome it is to work with a reasonable number of replicates, and it is almost impossible to perform this number of replications for each new compartment that occurs at the small scale. That is, while we can now derive very detailed information, the modeller and experimenter must jointly develop strategies to test representativeness or plausibility. From a modelling point of view, the need of accounting for what happens in the plant suggest that strategies to couple rhizosphere pore-scale models to single-cell models of the root physiology and development (e.g., Dupuy et al., 2008) should be investigated in the future.

Labelling experiments with radioactive ^11^C carbon (or ^14^C) provide information where recent assimilates are transported in relation to root type and position along the root (Schulte et al., in preparation; Holz et al., 2018), but such information is only available for distinct time points and for very young plants. Likewise, stable carbon isotopes (^13^C) can be used as a tracer for recent assimilates and the radial spread of ^13^C across the root tissue into the soil can be mapped in relation to tissue type and particle distribution, however, only for a small number of samples. Chemical quality of rhizodeposits can also be mapped and quantified with a resolution of 20 µm as recently shown for individual disaccharides (Lohse et al. submitted). With similar resolution, radial gradients of elements around roots can be mapped in soil using different microscopy techniques (Vetterlein et al. 2020). Only a few of these techniques, e.g., µXRF, have so far shown the potential to process a larger number of replicates (Lippold et al. in preparation). Methods to measure exudation rates from soil-grown plants (Oburger and Jones, 2018) are available and have great potential also for model parameterisation (see examples 3 and 4).

For structural gradients the available tools are already more powerful. With a spatial resolution in the range of 10-20 µm bulk density gradients around roots were investigated and statistically evaluated in relation to a range of different drivers (Phalempin et al. accepted).

It is a major advance that approaches have now been developed for chemical and structural parameters that truly measure the magnitude of radial gradients, in contrast to measurements in linearized (compartmentalized) or pseudo-linearized (rhizoboxes) systems. That we tend to see narrower zones with these methods (Lohse et al. submitted, Lippold et al. in preparation) is consistent with model predictions (Vetterlein et al. 2020, Fig. 4).

##### Microbiota in models

A very special case are the data on rhizosphere microbiota, the very tiny amounts of sample required for sequencing studies enables investigation of community composition down to the single aggregate level(Szoboszlay and Tebbe, 2021) and network analyses provides details on the interaction of the different species and how sensitively this is controlled by external drivers. Yet we lack quantitative data on functional properties or activities at the same resolution. There is a scarcity of data regarding the explicit distribution of microbes in general and of active ones in particular. Likewise, their habitat demands in relation to soil structure and resulting water, air filled pore space and carbon distribution can be addressed by modelling approaches, but validation of model output is challenging. Given the complexity and difficulty of obtaining non-destructive measures, confidence in models can be increased using simultaneously many measurable outputs related to the process(es) of interest as advocated by the so called pattern-oriented modelling approach (Grimm et al., 2005). In our opinion, validation using data from different levels of organisation such as emerging properties like respiration, change of C content and alike, or less available spatially explicit data (e.g., distribution of microorganisms on the root surface) can effectively test the assumptions included in the models, even when the data are not measured in the same experiment. Direct imaging of microbes in the pore space is possible with unspecific stains or Card-FISH, combined with stable isotope labelling, but the procedures are so tedious that it will be long until large datasets will be available. In view of the dynamic changes of microbiota along roots as they grow (Bonkowski et al., 2021) it is an open question whether focussing on specific microbes (CARD-FISH) or rather unspecific approaches are more promising for a given purpose. Each approach is able to tackle different questions. Targeted approaches can deal with biodiversity related questions or to focus on species with practical interest. An unspecific approach is more amenable to address question where the biodiversity is not the target but rather questions of CO_2_ release or total microbial biomass around roots.

##### The art of choosing the right drivers

Allowing for a few drivers only (in the priority program texture and genotype) bears the risk that important ones are missed out, or the chosen ones prove to be of no importance for the parameter in question. It turned out that both the soil- and the plant-oriented driver led to very interesting, sometimes surprising results. Two such opposing drivers have also served the function of motivating representatives of different disciplines to work together: i.e. plant scientists and microbiologist realized that their results are strongly dependent on texture, while physicists were challenged by the fact that plant’s feedback mechanisms are so smart that irrespective of substrate properties resources were exploited completely (Jorda et al. Kirk, Guy aration, Vetterlein et al. in preparation).

#### Further challenges and path ahead

Linking between more than two spatial scales or processes is still a challenge. Linking multiple scales and processes is the logical next step for future research and needs the combined use of experimental and modelling approaches.

So far, the impact of rhizosphere properties and processes have been discussed from the soil perspective. Except for water flow, a link between processes within the plant and how these react to or are coordinated with rhizosphere processes is still missing. Understanding which within-plant mechanisms control C-exudation is important to understand interactions between exudation and rhizosphere conditions. How local conditions influence growth such as mechanical rhizosphere properties but also transport properties that influence transport of signalling substances, e.g. ethylene, and thereby growth are examples of two-way feedbacks between growth and rhizosphere properties and processes.

Another aspect is the temporal scale of the rhizosphere organisation. In this paper (and the priority program), we focussed on the dynamics around a growing root system of an annual plant (maize). Whether such a rhizosphere system has a legacy after the plant died off that is of benefit for subsequent generations or whether it rather has negative effects requires further research. This would call for investigating multi-annual self-organisation effects of the rhizosphere in annual cropping systems. Understanding these effects is important to design and manage crop rotations and no-till systems.

## Declarations

### Availability of data and material (data transparency)

All data are available on request from the authors.

### Code availability

Code is available on request from the authors.

### Authors’ contributions

Our opinion is the result of many discussions we have had during meetings and workshops within the framework of the German priority program “Rhizosphere Spatiotemporal Organisation – a Key to Rhizosphere Functions”.

### Additional declarations for articles in life science journals that report the results of studies involving humans and/or animals

Not applicable

### Ethics approval

Not applicable

### Consent to participate

Not applicable

### Consent for publication

All authors agreed with the content gave consent to submit this manuscript.

