## SupplementaryInformation for "Linking rhizosphere processes across scales: Opinion"

**Additional information to the experiment described in example 1**

**Plant Growth**

Maize plants (*Zea mays L.*) were grown in 3D printed seedling holder microcosms each consisting of a seed compartment and seven syringe barrels of 1ml volume, following the approach of Keyes et al. (2013) and Koebernick et al. (2017). The seedling holders were filled with a loamy substrate (haplic Phaeozem) which was sieved and fertilized according to the protocol of Vetterlein et al. (2021). The seeds were pre-germinated on filter paper for 48 hours before planting them individually into the microcosms. The plants were grown in a climate chamber for 14 days under the conditions defined by Vetterlein et al. (2021) before they were transported to the TOMCAT facility at PSI, Switzerland. To enhance the contrast between different materials in the CTs, samples were dried down via transpiration prior to the harvest and scanning procedure.

**Imaging, Image Processing and Mesh Generation**

In situ measurements were performed at the TOMCAT beamline of Swiss Light Source (SLS). At a beam energy of 20keV, a set of three syringe barrels per seedling holder was scanned at three different positions each - at the centre ±1.5cm. The resulting local tomographies of 1.4mm height comprised 3501 projections with an exposure time of 80ms per projection. The data were collected using a detection system comprising of a C20-72 Scintillator, an Optique Peter Microscope with a magnification factor of 10 and a pco.Edge 5.5 detector. This setup allowed for an actual pixel size of 0.65µm. Regarding raw data reconstruction and filtering the Gridrec algorithm (Dowd et al., 1999; Marone and Stampanoni, 2012) as well as the Paganin filter (Paganin et al., 2002) were applied.

Image stacks comprising of 2160 slices per sample were processed in Avizo 2020.1 (Thermo Fisher Scientific). Conversion to 8-bit grayscale images as well as extracting regions of interest including all root hairs led to a reduction in computation time for later steps. Sharpening of the images was achieved by the “Unsharp Masking” filter before segmenting the different domains (soil, air, and root) by applying a watershed transformation. Morphological opening and closing algorithms were used to separate roots from hairs and to remove laterals. A label analysis was performed to measure the surface area of the epidermis and the porosity of the samples. The soil domain was smoothed by the “Binary Smoothing” module. Avizo’s “Generate Surface” module was utilized to obtain tetrahedral surface grids of the soil, root, and hair domains, which were simplified within the “Simplification Editor”. The grid of the soil domain as well as the grid representing the contact between soil and root (± hairs) were exported individually as stl-files and imported into Gmsh (Geuzaine and Remacle, 2009), where they were merged to a watertight volume. Physical tags for incorporating boundary conditions were defined and meshing was conducted using the frontal algorithm (Rebay, 1993). The quality of the 3d meshes was optimized by applying the Netgen optimizer in Gmsh (Schöberl, 1997) and the exported msh-files were converted to the foam format.

**Simulations and Post-Processing**

Time stepping was accomplished by the backward Euler method and the Gauss scheme was used for discretizing the Laplacian term. Interpolation scheme was set to “linear” and surface normal gradient scheme to “corrected”. A total diffusion time of 1h per case was simulated using a Fujitsu Celsius R970power workstation.

Simulation results were post-processed using the softwares Paraview (Ahrens *et al.*, 2005) and RStudio (version 1.2.1335, RStudio Team, 2020). The total amount of diffused carbon into soil was calculated by integration of the concentration data over the volume of the soil domain utilizing the “Integrate Variables” filter of Paraview. Carbon concentration in relation to the distance from the root-surface was computed in Paraview and plotted in RStudio.
